# *The Incubot*: A 3D Printer-Based Microscope for Long-Term Live Cell Imaging within a Tissue Culture Incubator

**DOI:** 10.1101/2020.05.28.121608

**Authors:** George O. T. Merces, Conor Kennedy, Blanca Lenoci, Emmanuel G. Reynaud, Niamh Burke, Mark Pickering

## Abstract

Commercial live cell imaging systems represent a large financial burden to research groups, while current open source incubator microscopy systems lack adaptability and are sometimes inadequate for complex imaging experimentation. We present here a low-cost microscope designed for inclusion within a conventional tissue culture incubator. The build is constructed using an entry level 3D printer as the basis for the motion control system, with Raspberry Pi imaging and software integration, allowing for reflected, oblique, and fluorescence imaging of live cell monolayers. The open source nature of the design is aimed to facilitate adaptation by both the community at large and by individual researchers/groups. The development of an adaptable and easy-to-use graphic user interface (GUI) allows for the scientist to be at the core of experimental design. Simple modifications of the base GUI code, or generation of an entirely purpose-built script, will allow microscopists to place their experimental design as the priority, as opposed to designing experiments to fit their current equipment. The build can be constructed for a cost of roughly C1000 and thus serves as a low-cost and adaptable addition to the open source microscopy community.

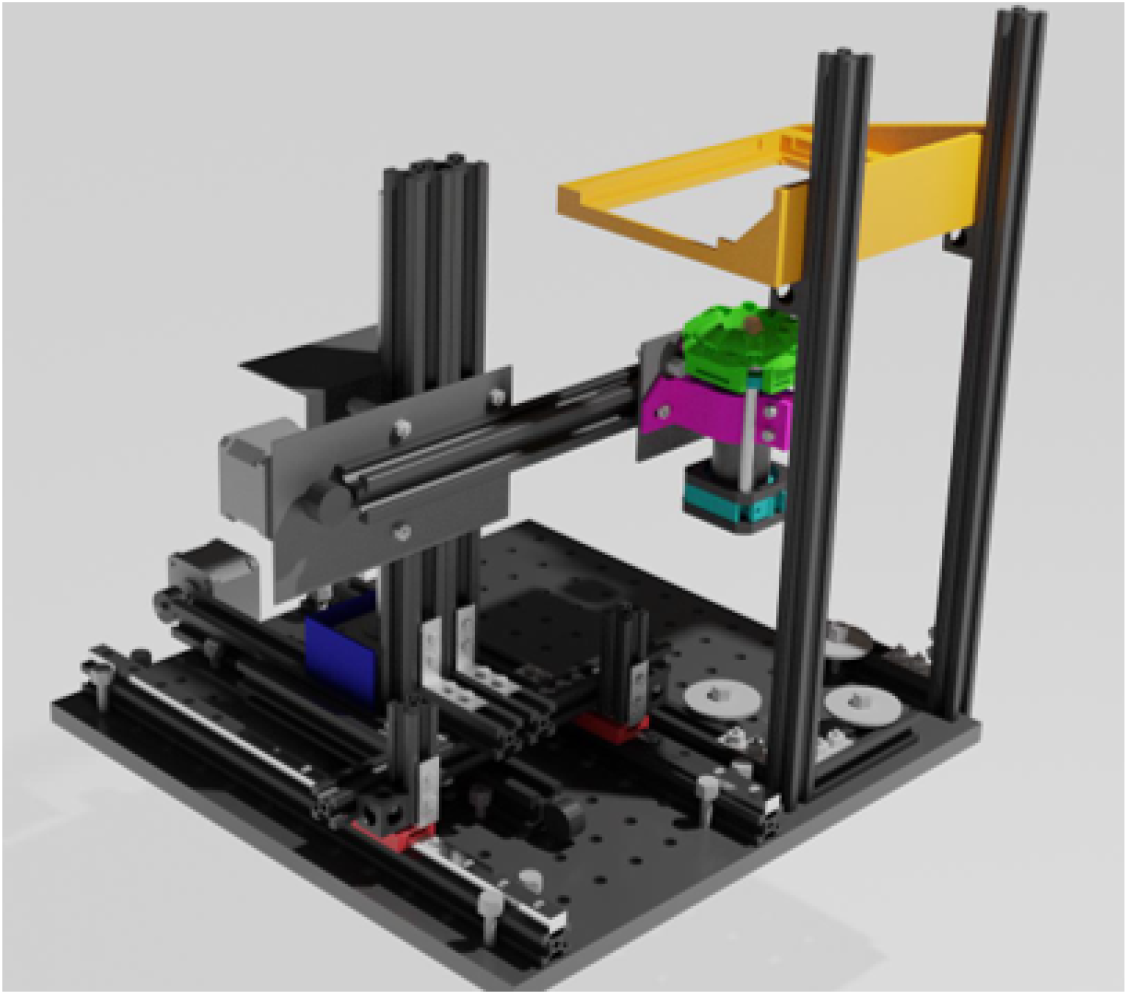

## Hardware in Context

Live imaging of cells under physiological conditions is an essential technique in biomedical sciences for analyses of cell proliferation (1,2), cell migration (3,4), and cell-cell interactions (5). Two commercial options are available to researchers aiming to perform live-cell imaging. First, purchase a specific stage incubator designed to upgrade the current expensive microscopy system you already own. Alternatively, purchase a whole new system designed only to be use within a tissue culture incubator, often with a significant price tag, with additional costs for any add-ons such as “automated scanning”. Despite the widespread use of live imaging, the cost of the equipment necessary for such types of experiments remain high. A common way to deal with the significant cost is to arrange a consortium or multi-group effort to purchase the equipment, sharing the resource after purchase, or institutional investment in core imaging facilities. In the case of shared equipment, long-term studies can prove difficult to schedule and can reduce flexibility in experimental design, particularly in early stage, exploratory and scoping studies. Usage fees applied to maintain the equipment may also then limit access to lower income laboratories. This cost barrier to owning complex imaging equipment can stifle research efforts in the biomedical field.

The open source movement within science has been providing innovations in affordable laboratory equipment, including microscopy and optics (6–9). Open source approaches for low-cost incubator based microscopes have been explored (10–12) and serve as great examples of affordable microscopes for live-cell imaging. However, several limitations with current designs curtail their potential widespread use within laboratories. The use of a CNC machine is neither common nor accessible amongst biomedical researchers. Options that allow for in-incubator microscopy but lack a motion control system result in a low-throughput. The use of only transmitted white light reduces the range of potential applications of such builds. An ideal scenario would be an affordable open source system that makes use of conventional low-cost commercial products while allowing for adaptability as a group’s research interests or capabilities may change.

Microscopy is always a trade-off between field of view (FOV) and resolution. A static microscope is simple and effective, but only allows a small FOV to be imaged. Imaging more than a single FOV requires some element of motion control of wither the optics or the sample. The open flexure microscope is a remarkable solution to this problem, as it includes very precise motion control at a low cost (13). The main drawback is that the range of movement is limited and do not allow for scanning plates or flasks that are routinely used in cell culture experiments. Luckily, the problem of precise and repeatable control of three-dimensional movement has been solved in another field: 3D printing. Commercial 3D printers require sub-millimeter precision over a range of travel of tens of centimeters.

Therefore, we designed an incubator microscope, the *Incubot*, which repurposes the core motion control system from a simple, entry-level cartesian 3D printer (a Tronxy X1). An associated easy-read assembly guide has been made available on the online repository to help explain all steps in construction of this system, with the aim that non-experts in microscopy or optics can construct and use this equipment.

## Hardware Description

The *Incubot* uses a Tronxy X1 3D printer as a motorised stage for optics movement allowing for XYZ imaging of tissue culture flasks or plates (Fig 1.). The X and Z axes are coupled directly to the Y axis to allow for the 3-dimensional movement of the optics unit which is coupled to the extruder plate. A tissue culture plate is maintained above the optics via a stage composed of 15×15 mm aluminium extrusion and 3D-printed components, which remains stationary while the optics are translated. Moving optics under a static sample prevents movement of the medium during imaging, which may affect the cells. The optics unit is composed of 1” ø tubes, connecting a commercial infinite-conjugate objective lens to a Raspberry Pi Camera (V2), using an achromatic doublet tube lens to allow for microscopic image acquisition. The optional inclusion of a long pass filter allows for fluorescent imaging of samples. A silicone moulded rig was used to mount 8 LEDs around the objective lens, and thus allow for several illumination techniques to be performed, such as reflected, o blique, a nd fluorescent. Bo th th e motorised stage and the plate-holder are mounted onto a 300 mm × 300 mm breadboard (M6) with vibration-absorbing feet. The whole unit can be placed within a tissue culture incubator for long term use. This setup allows for inverted imaging of a tissue culture plate/flask within an incubator maintained at physiological conditions and avoids the negative impact of thermal drift associated with heated stages.

**Figure 1.**
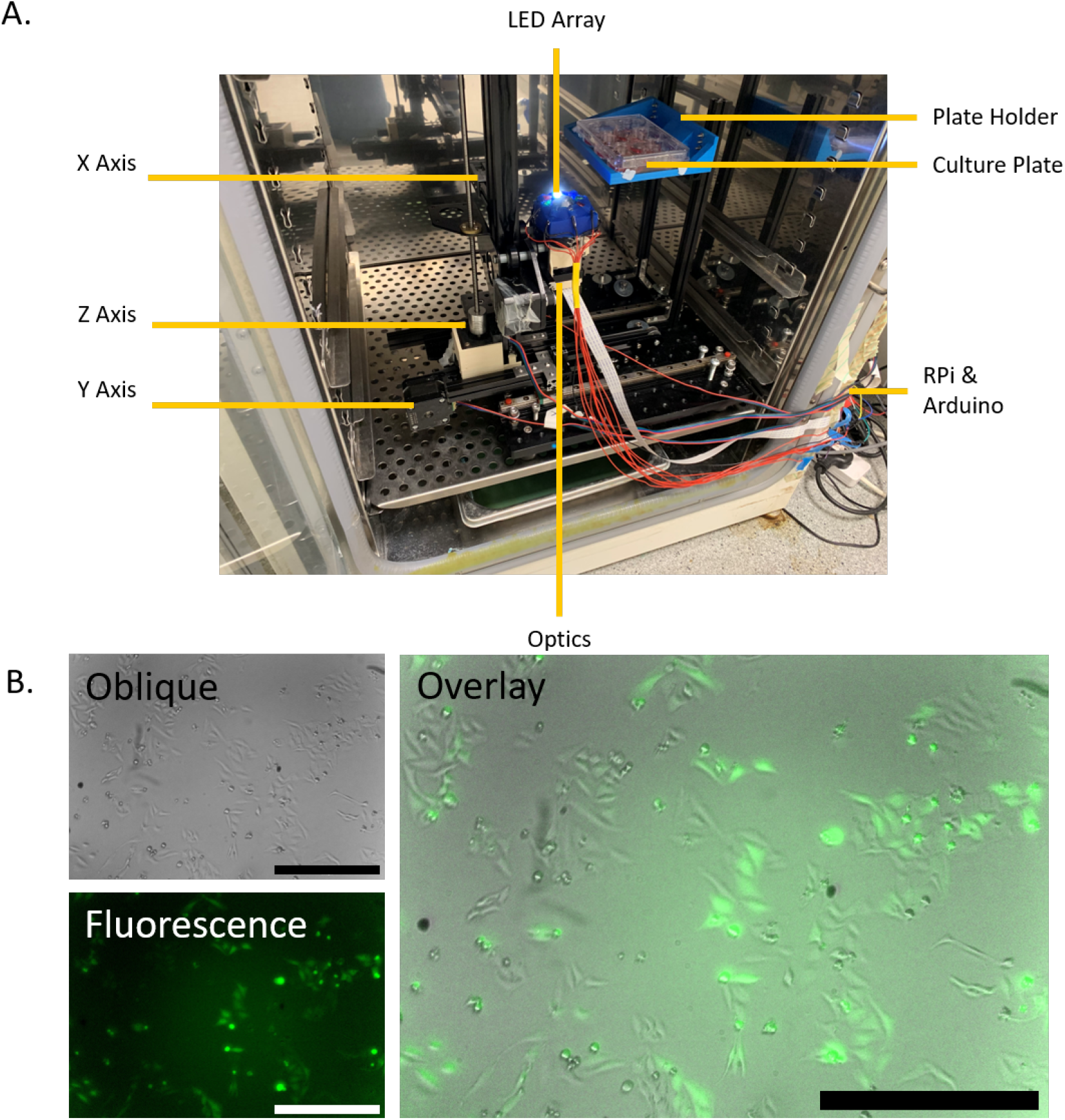
A) Fully Assembled *Incubot* within a Tissue Culture Incubator with Key Components/Features Annotated. B) Representative images of HeLa-GFP live cells within an incubator using the *Incubot* using oblique white illumination (Oblique), blue LED excitation of GFP (Fluorescence) and the two images overlain (Overlay). Scale bar indicates 500 μm.

A Graphical User Interface (GUI) was developed using Kivy for Python 3, allowing for simple user control over parameters for live cell imaging, including the number of time points to image over, the temporal spacing between them, the number and layout of wells, the area of each well imaged, and automated focusing for each well. The total cost of this system (€C1061.79) represents a substantial saving over commercial equipment (14).

This build represents a low-cost alternative to commercial microscopy systems for physiological imaging of cells, with specific utility in simple imaging experimentations. The use of simple components and 3D printing for construction of the build allow for modification and adaptation of the build for individual users based on their needs.

Potential uses of the *Incubot*:

- Long-term monitoring of tissue culture experiments
- Long-term monitoring of tissue culture flasks for determination of optimum time for passaging/experimentation
- Semi-portable device for tissue culture information outreach
- Use as a regular scanning microscope for fixed cell imaging

## Design Files Summary

All files relating to the *Incubot* can be found at our public online repository: https://osf.io/es3hr/

### XYZ_Coupler

3D printed unit designed to couple the X and Z axes of the original Tronxy X1 to its Y axis.

### Optics_Holder

3D printed unit to replace the extruder portion of the original Tronxy X1. Used to clamp hold of the optics and to couple XYZ motion to the camera.

### Pi_Camera_Holder

3D printed unit to hold the Raspberry Pi Camera in the correct orientation and distance from the lenses of the optical system.

### LED_Holder_Mould

3D printed negative mould for casting a silicone LED-holding ring.

### Plate_Holder

3D printed unit to allow for stable elevation of a tissue culture plate over the optics of the *Incubot* for imaging.

### MotionValidationProtocol

Python script to perform sequential positive and negative movements of known distance in the X and Y axes independently. Images are acquired after each movement to allow for later confirmation of movement distances.

### IncubotGUI

Python script that uses Kivy to generate a graphical user interface for interacting with the *Incubot* and performing imaging experiments.

### StationaryStabilityTesting

Python script to perform repeat imaging of a stationary location over time for XY drift tracking.

### MotionRestTesting

Python script to determine the necessary time between movement commands and image acquisition for optimum *Incubot* function.

### PairwiseStitchingMotionValidation

ImageJ Macro to stitch sequential images from the Motion-ValidationProtocol.py script for analysis of motion distances moved.

### Movement Validation Template

Excel file template to input data resulting from the PairwiseStitchingMotionValidation.ijm ImageJ macro.

## Build Instructions

### General Safety Notice

The build will ultimately require a 12V power supply to the CNC motor shield, and 5V power supply to the Raspberry Pi. Take care when performing any soldering to prevent short circuit generation or injury while working with intense heat.

### General Assembly Notice

There are many connections listed below that rely on a M3 screw and a M3 nut. Please note that in most cases it is possible to reverse the orientation of these while constructing the *Incubot*, with no negative impact on build stability or quality. If you find inserting the screws into a MakerBeam groove to be easier during construction than sliding the nut, then please do whatever is easiest for you.

An animated assembly overview can be found within the OSF repository.

### Rail Assembly (Fig 2.)

1. Orientate the M6 breadboard (BB) on a flat surface.
2. Roughly align two 15×15×300 mm MakerBeam beams (MBXL300) on the BB 110 mm apart.
3. Incorporate 13 M3 nuts (M3N) into each MBXL300 topmost groove. Ensure these are evenly spaced and will line up with the holes in the 300 mm rails (MBR).
4. Align a MBR with each MBXL300, pressing the ends of both components against a flat surface. Ensure the M3N from step 1 are aligned with the holes in the MBR.
5. Insert a M3 6 mm screw (M3S6) into each of the holes of the rail, and loosely screw into the underlying M3Ns.
6. Once all M3S6s are loosely fit into their M3Ns, double-check the alignment of the MBR and the MBXL Alternating between all the M3S6s, gradually tighten until all M3S6s are strongly tightened, maintaining alignment throughout the tightening process.

- Aligning the MBR and the MBXL300 at the ends can be done by pressing the ends of both against a flat object and maintaining pressure during the screw tightening process.
7. Use large-head M6 screws (M6S20) to affix the rails to the BB. The large heads of the screws are to be used as clamps to secure the rail units to the BB. Use two screws at each end of each rail unit, as close to the end as possible so as not to impede movement along the rail carriage.
8. Ensure the rail units are affixed with a separation of 110 mm and are maintained parallel along their axes.

**Figure 2.**
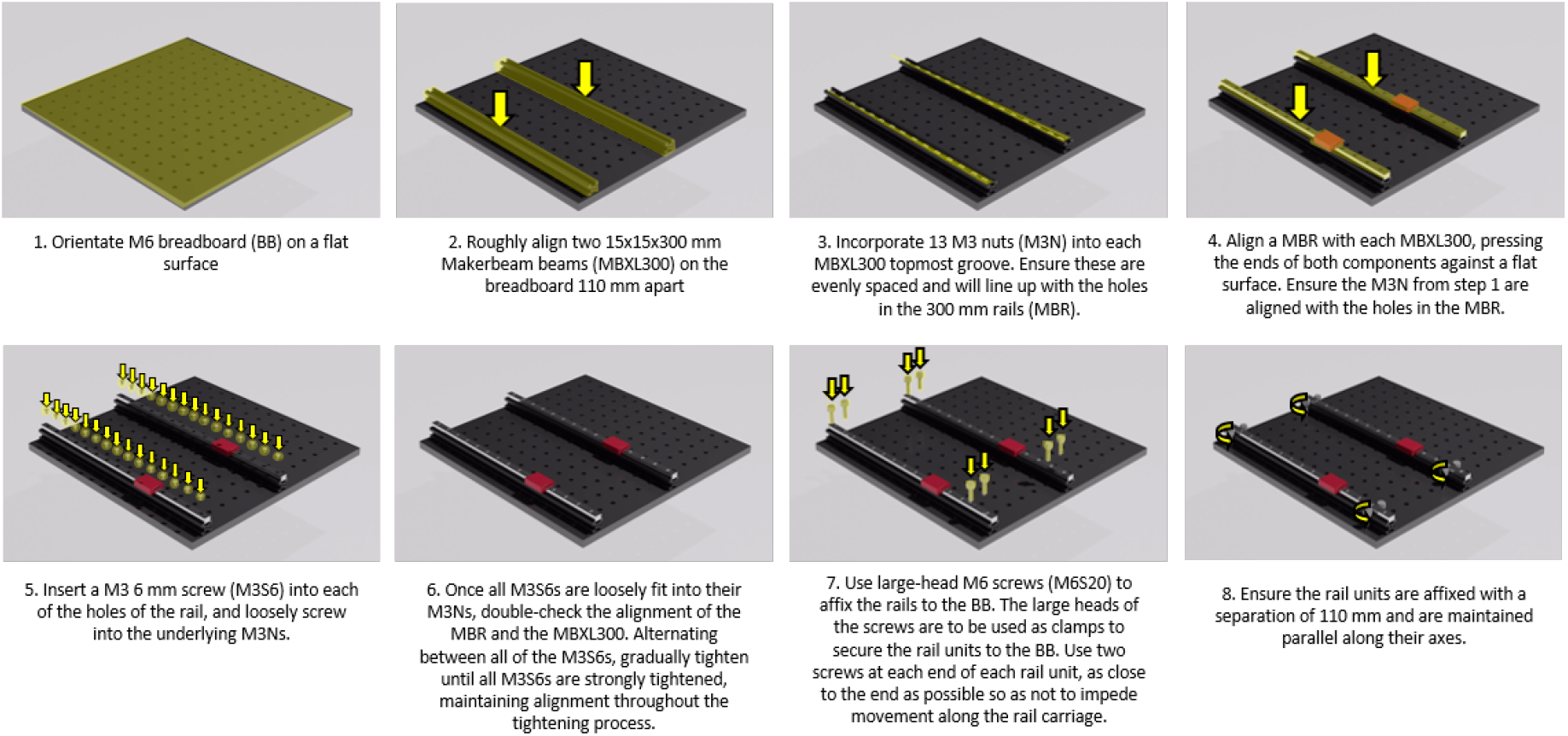
*Incubot* Assembly Visual Guide – Rail Assembly

### Coupling the X, Y, and Z Axes (Fig 3. 4.)

1. Affix a corner bracket (MBCB) to each of the rail carriages using two M3S6s.
2. Align a 50 mm MakerBeam XL beam (MBXL50) with each of the MBCB and insert 2 M3Ns into each in the groove facing the MBCB.
3. Align the M3Ns with the holes in the MBCBs and affix to the MBCB using a M3S6 for each M3N.
4. Insert an additional M3N into each MBXL50 into the groove facing away from the other MBXL50, leaving it to rest at the bottom of the MBXL50 groove.
5. Insert 4 large-head M6 screws (M6S20) into the BB with roughly 9 mm of threading left above the BB.

- Orientate these screws so that there is one pair in the very back row of BB holes, and the other pair is roughly 230 mm away. The screws within each pair should be in adjacent BB holes.
6. Attach the wheels to the Y axis of the Tronxy X1 3D printer (X1P) using M5 50 mm screws (M5S50) and the 3D-printed wheel spacer (3DWS).

- This unit will not be firmly adhered as the nuts (M5Ns, originally from X1P) will not be attached until later, so be careful when moving this unit until then.
7. Insert a M3 12 mm screw (M3S12) into each of the two holes at the front end of the X1P plate, and loosely affix a M3N to each of the M3S12s achieving a loose fit where the screw head is partially mobile.
8. Slot a MakerBeam XL 200 mm beam (MBXL200) onto the X1P plate surface, aligning the M3S12 with its bottom-most groove. Tighten the M3Ns from below until firmly affixed, ensuring the MBXL200 is centred relative to the X1P plate.
9. Slide the Y axis of the X1P onto the BB so that the M6S20s (step 5) are resting in the large groove on each side of the axis.
10. Tighten the M6S20s in the BB (step 5) until the Y axis is firmly adhered to the BB, ensuring the X1P plate is allowed the full range of motion possible, and attach the X1P plate through the M5S50 screws used in wheel incorporation.
11. Place 2 M3Ns in the back groove of the MBXL200, bringing close to the centre.
12. Place 2 M3Ns in the front groove of the MBXL200, one at each end, leaving it lateral to the MBXL50.
13. Align a MakerBeam XL corner cube (MBXLCC) with each MBXL50 at the junction between the MBXL50 and the MBXL200 with the nuts aligned to the holes of the MBXLCC.
14. Affix each MBXLCC using these M3Ns and M3S6s to couple the MBXL50 to the MBXL200.
15. Gather 2 new MBXL50s and insert a M3N within the bottom groove of each. Align a MBXLCC with each M3N, and affix using a M3S6, ensuring alignment with the end of each MBXL50, forming a “L” shape.
16. Rest the unit from step 15 on the MBXL200 top surface, so that the angle created by the MBXLCC addition fits perfectly onto the corner of the MBXL200.
17. Insert a M3S6 into the build plate from below and attach a M3N to hold it loosely in place.
18. Slide a MakerBeam XL 150 mm (MBXL150) and a MakerBeam XL 100 mm (MBXL100) through the right and left M3Ns (step 17) of the X1P plate, respectively. Tighten the screws.

- Ensure the shorter beam is on the same side as the Y axis motor to prevent obstruction of the full Y axis movement.
19. Insert 2 M3Ns into the top groove of each of the MBXL units from step 18, and 6 M3Ns into the top groove of the MBXL200, with 3 M3Ns on the left, and 3 M3Ns on the right.
20. Rest a MakerBeam XL T-bracket (MBXLTB) on the junction between the MBXL200 rod and the MBXL150 unit. Align the M3Ns with the holes in the MBXLTB.
21. Affix the MBXLTBs to the MBXL beams using M3S6s through each of the M3Ns.
22. Affix the MBXL50/MBXLCC units (step 13) to the back groove of the MBXL200 using two M3S6s, one for each of the M3Ns placed in the back groove of the MBXL200 (step 12).

- These will be used to couple the XZ axes, so ensure they are in the correct position. To orientate these correctly, it may be necessary to perform steps 23 and 25 (not step 24) and mark the locations of the front groove of the Z axis on the MBXL200.
23. Place the 3D printed XYZ Coupler (3DXYZC) on the X1P plate through the inverted M5S50s used for wheel incorporation.
24. Affix the 3DXYZC unit to the plate using M5Ns (originally from X1P).
25. Insert the X1P Z axis bottom end into the front portion of the 3DXYZC, ensuring the Z axis stepper motor is resting on the back portion of the 3DXYZC.

- You may need to loosen and then tighten the Z axis motor from the Z axis beam to align this properly.
26. Insert 2 M3 T-nuts (M3NTs) into each front groove of the Z axis beams.

- This step cannot support the swapping of the nut with the screw, as the M3NTs have the required dimensions to support coupling.
27. Insert 2 M3Ns into the top groove of each of the MBXL50s (step 15).
28. Orientate a MBCB at the junction of the MBXL50 beams and the Z axis beams.
29. Loosely affix the MBCB to the MBXL50 beams using 4 M3S6s, one for each M3N.

- Leave loose enough to be able to move the MBCB along the MBXL50 beams
30. Use 4 M3S12s to affix the MBCB to the Z axis beams through the M3NTs, ensuring the units rotate in a way to prevent their exit from the groove.

- To do this, ensure the screw is in the threading of the M3NT, then use the screw to orientate the M3NT so that its long angle is perpendicular to the groove opening. Use the screw to pull the M3NT so that it is pressed against the groove, and then screw while maintaining the tension so the M3NT units maintain their orientation.
- It may be necessary to establish the M3S12 with the M3NT through the MBCB before inserting the M3NT into the Z axis groove.
31. Tighten all screws from steps 29 until firmly attached while maintaining alignment. The X, Y, and Z axes are now fully coupled to each other in addition to the Y axis support rails.

**Figure 3.**
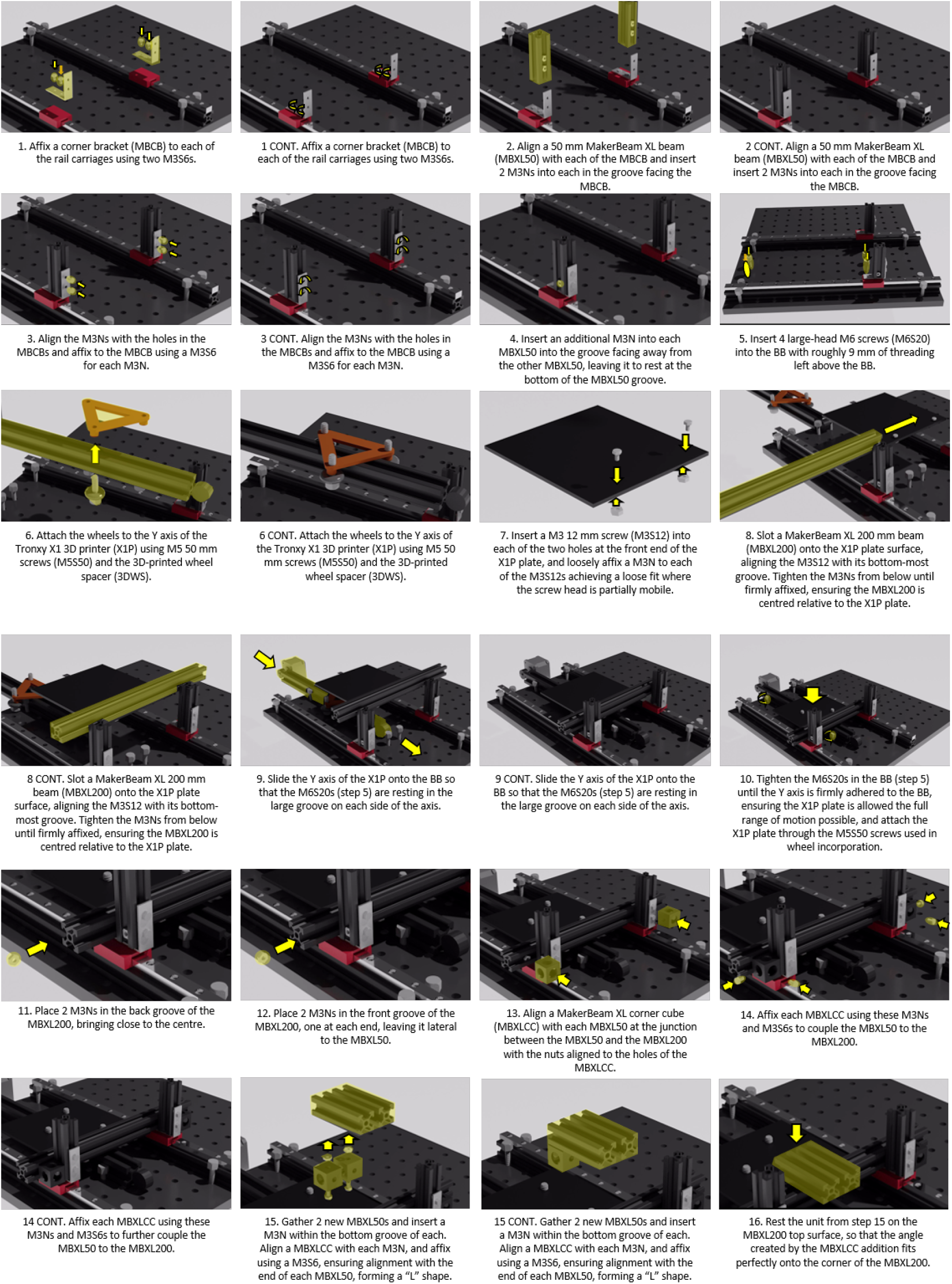
*Incubot* Assembly Visual Guide – Axes Coupling Part I

**Figure 4.**
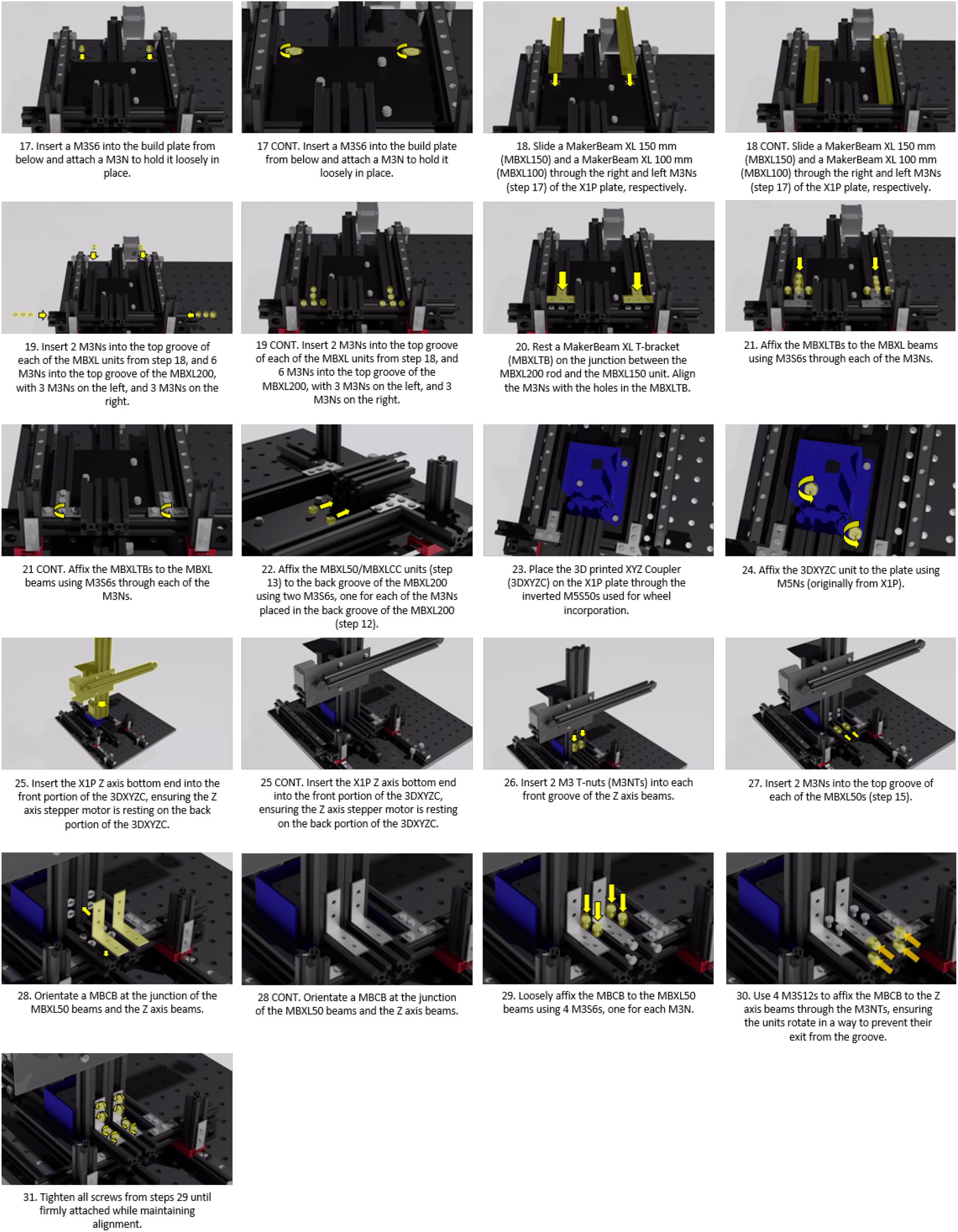
*Incubot* Assembly Visual Guide – Axes Coupling Part II

## Optics Assembly (Fig 5.)

1. Construct the Optical Pathway

a. Orientate a lens holding ring (LHR) within the half-inch SM1 tube (SM1L05) at the male end of the tube.
b. Carefully place the tube lens (TL) in the SM1L05 tube with the convex surface facing the female end.
c. Secure the TL in place within the SM1L05 using a second LHR.
d. Screw the male end of the SM1L05 into the female end of a one-inch SM1 tube (SM1L10) until fully tightened.

- If fluorescent capabilities are required, incorporate the long-pass filter (LPF) into the SM1L10 using an LHR before attaching to the SM1L05
e. Cover both ends of this tube with parafilm until later use to prevent dust settling on the tube lens.
2. Construct the Camera Unit

a. Remove the lens from the Raspberry Pi Camera V2 (PiCV2).

- While it is possible to use pliers for this, several open source 3D printed options exist which reduce the risk of scratching the sensor and damaging the camera
b. Attach the long ribbon cable (RCL) to the PiCV2.
c. Place the PiCV2 into the appropriate depression within the Raspberry Pi Camera-holder (PiCH).
d. Hold the PiCV2 firmly within the depression using a Thorlabs cage plate (TLCP).
e. Place an additional TLCP on the other side of the PiCH, maintaining pressure on the PiCV2 from both sides.
f. Insert two 1” cage assembly rods (CAR1) through opposite corner holes to align the plates, and thus the PiCV2.

- One rod should be to the left of the RCL when the PiCV2 is orientated with the lens facing upwards.
g. Tighten the mounting screws on CAR1 units to secure the camera unit in place.
3. Screw the entire unit resulting from step 1 into the camera unit generated in step 2.
4. Attach the SM1 to RMS adapter (SRA) to the lens tube and tighten until fully secured.
5. Screw the objective lens (OBJL) into the SRA until fully tightened.
6. Construct the LED Holder

a. Generate 50 g of Dragon Skin silicone (DSSil) by mixing 25g of part A with 25g of part B and mixing thoroughly. Cast the DSSil to the LEDHM.

- It may be necessary to vacuum the silicone to remove large air bubbles, however this is unlikely to be necessary unless you are overly enthusiastic with your mixing.
b. Seal the chamber with the lid and cure (either room temp overnight or higher temp for faster curing).
c. Remove the DSSil from the LEDHM once fully cured to get the silicone LED holder (SilLEDH).

- Use tools to try and pry free the lid portion from the mould. This is difficult and may break the lid of the LEDHM in the process.
d. Insert LEDs in the appropriately sized holes within the SilLEDH, forcing their pins through the silicone. Arrange the LEDs in the pattern: White, Red, Blue, Green, White, Red, Blue, Green. This will lead to pairs of the same colour LEDs opposite each other (Fig 6.).

- Each LED should be orientated at 45° to the place of the SilLEDH.
e. Solder the LEDs to wires on their positive and negative pins.
f. Collate the ends of the negative wires onto a single small metal ring and solder to secure.
g. Solder a long wire to this metal ring, and then insert the free end into a female-female jumper wire.
h. Label each of the positive wires appropriately.

**Figure 5.**
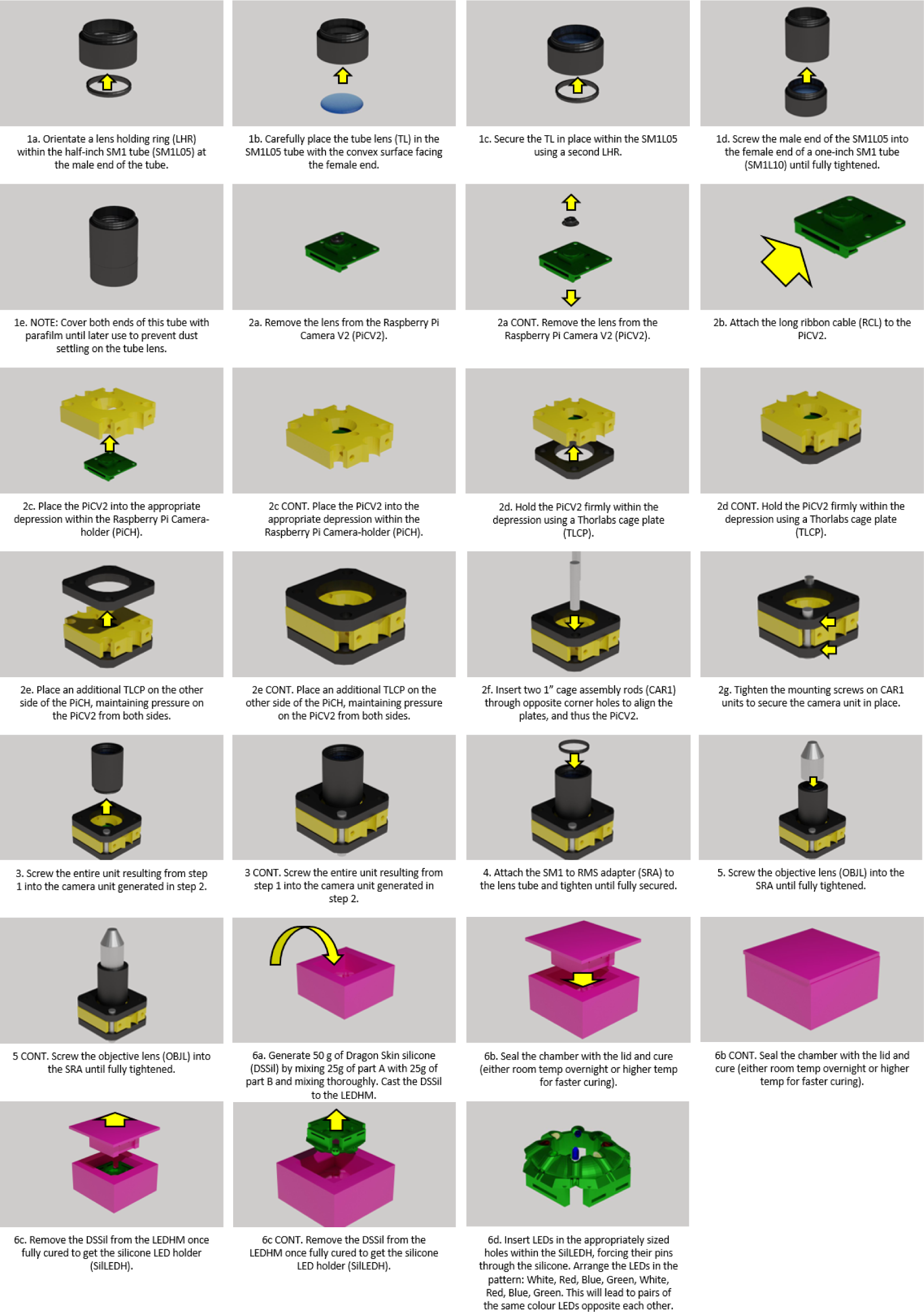
*Incubot* Assembly Visual Guide – Optics Assembly

**Figure 6.**
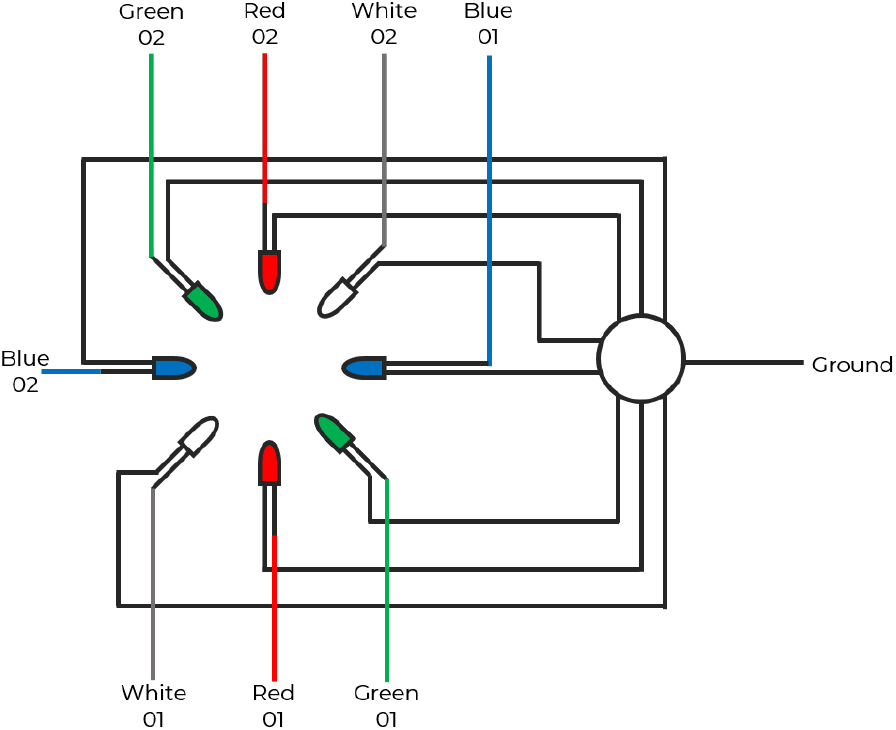
*Incubot* Assembly Visual Guide – LED Arrangement Schematic

## Optics Coupling (Fig 7.)

1. Orientate the 3D printed optics holder (3DOH) with the extruder plate using M3S12 units.
2. Attach the 3DOH to the plate using M3Ns and firmly tighten the M3Ns until the 3DOH is stable against the extruder plate.
3. Stabilise the extruder plate on the X axis beam using the wheels, M5Ss, spacers, and M5Ns (all originally from the X1P).
4. Insert a M3N into each of the hex spaces in the tightening portion of the 3DOH.
5. Insert a M3 25 mm screw (M3S25) into each of the tightening spaces on the opposite side to the M3Ns and loosely tighten.
6. Insert the optics unit into the hole in the 3DOH and tighten the M3S25 until the optics unit is held in place by the tension of the 3DOH.
7. Insert the 3” cage assembly rod (CAR3) into the corner hole of the top TLCP and tighten the cage assembly screw.
8. Attach an additional TLCP onto the top of the CAR3 and tighten its cage assembly screw to hold in place.
9. Stretch the constructed silicone SilLEDH over the top TLCP so that it is securely affixed to the optics unit, and the LEDs are surrounding the OBJL at an appropriate level for illumination of a sample without impeding the focal length of the OBJL.

- Confirm a roughly 45° angle between the objective direction and the LED orientation to allow for oblique imaging later.

**Figure 7.**
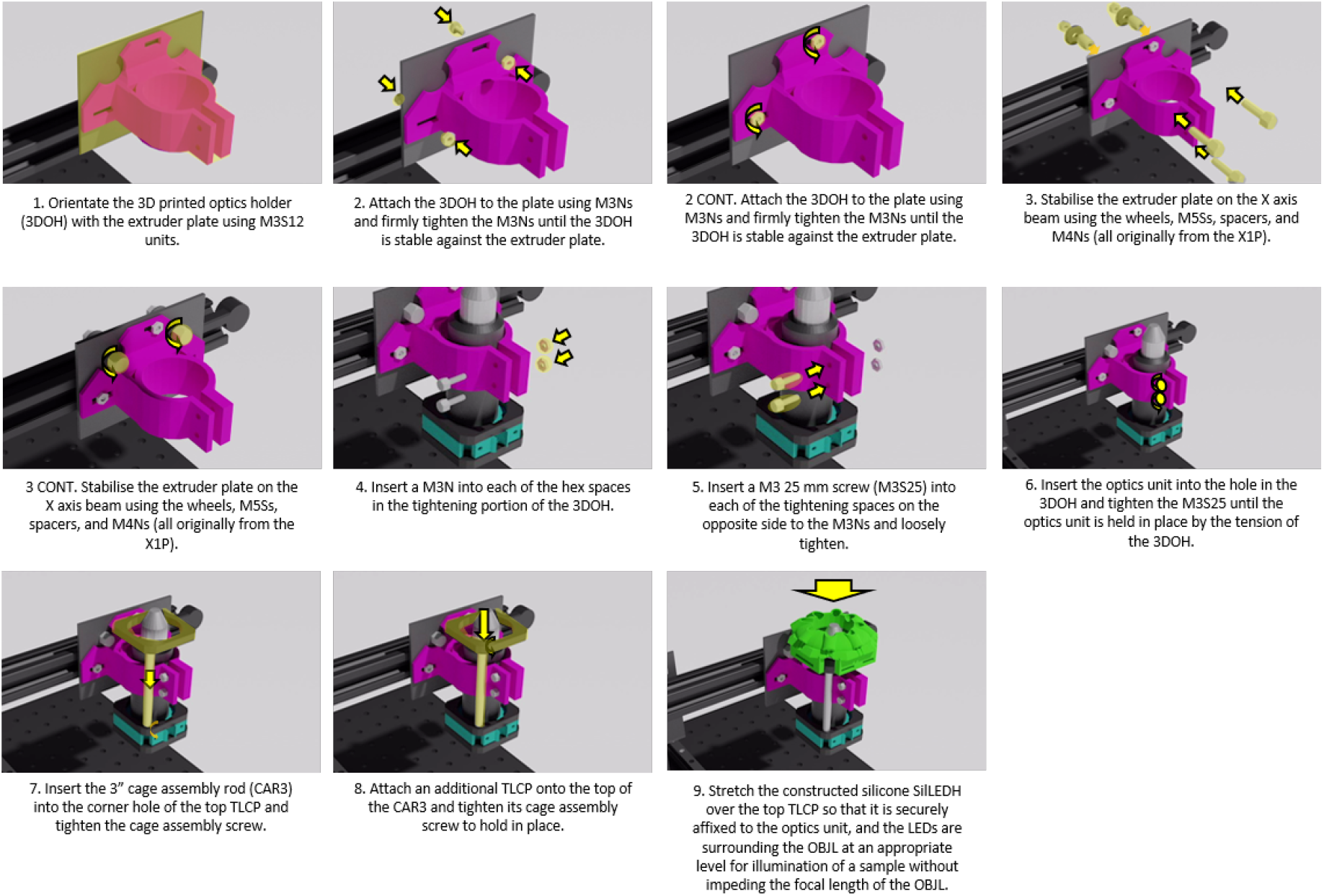
*Incubot* Assembly Visual Guide – Optics Coupling

## Plate Holder Construction (Fig 8.)

1. Slide a M3S6 unit into a groove on a MBXL300. Repeat for a second MBXL300.
2. Incorporate a MBXLCC into each of the M3S6 units.
3. Affix to the MBXL300 using a M3N, ensuring the MBXLCC units are level with one another, and that they are roughly 80 mm from the top end of the MBXL300.
4. Incorporate 3 M3S12 units in the smaller portion of the groove on each MBXL300 formed by the MBXLCC.
5. Confirm the M3 holes within the 3D printed plate holder (3DPH) are large enough for M3 screws to fit in without threading.

- It may be necessary to use a Dremel or alternative to enlarge the holes, depending on the tolerances of your printer.
6. Slide the 3DPH onto the exposed M3S12 units in each of the holes on its back surface.
7. Use a M3N on the top and bottom of each of these sets of screws and loosely tighten.
8. Ensure the bottom of the 3DPH is pressed firmly against the top of the MBXLCC units, and that the MBXL300 units are running perfectly in parallel.
9. While maintaining alignment and organisation, tighten the M3Ns in sequence until the 3DPH is firmly attached to the MBXL300 units.
10. Attach a MBCB to a MBXL50 unit using 2 M3S6 units, and 2 M3N units.
11. Repeat step 10 two times to get a total of three units.
12. Attach these units (steps 10 11) to the base of the MBXL300 units from step 9 using two M3S6s and 2 M3Ns for each MBCB. Attach 2 of the units to the groove on the side of the 3DPH, and the 3rd unit to the medial groove of the right MBXL300.
13. Incorporate the plate-holding unit onto the BB using M6S25s. It may be necessary to apply a large washer to improve stability of the plate-holder relative to the BB.

**Figure 8.**
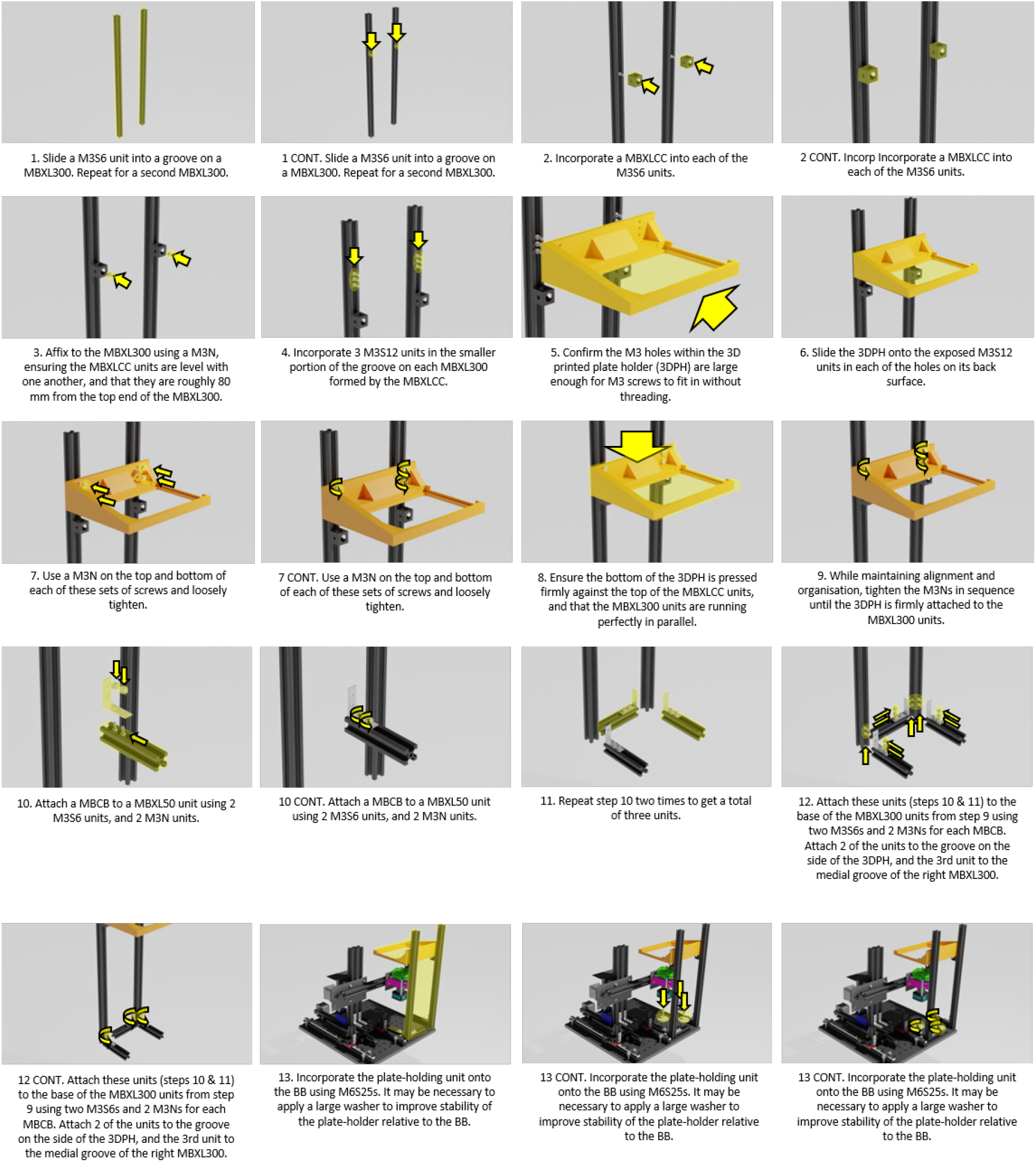
*Incubot* Assembly Visual Guide – Plate Holder Construction and Coupling

## Electronics (Fig 9.)

For a full guide on CNC motor shield establishment, please see (15).

1. Affix the heat sinks to the stepper motor drivers (SMD) (16).
2. Apply any connectors to the microstepping pins on the CNC motor shield (CNCMS) that you wish, keeping note of the impact this will have on microstepping and torque ability.
3. Insert the SMDs into the appropriate positions for the X, Y, and Z axes of the CNCMS, leaving the A axis without a driver and ensuring the SMDs are orientated correctly.
4. Attach a red wire to the positive inlet of the CNCMS, and a black wire to the negative inlet of the CNCMS.
5. Solder/connect these wires to the corresponding wires/connectors of a 12 V/1250 mA power supply.
6. Connect the stepper motor cables to their appropriate pins on the CNCMS: X-X, Y-Y, Z-Z, leaving the A pins blank.

- Depending on the distance from your motors to your CNC motor shield, it is likely necessary to replace the cables that come standard with the X1P with a longer alternative.
7. Attach the CNCMS to the Arduino Uno R3 (AUR3), ensuring all pins are firmly in their appropriate openings, taking care not to bend or damage any pins.
8. Connect the AUR3 to the Pi3B+ using a USB cable.
9. Establish the GPIO pins and any additional cables as specified (Fig 9.).
10. Connect the AUR3 to the Pi3B+ using the USB cable.

**Figure 9.**
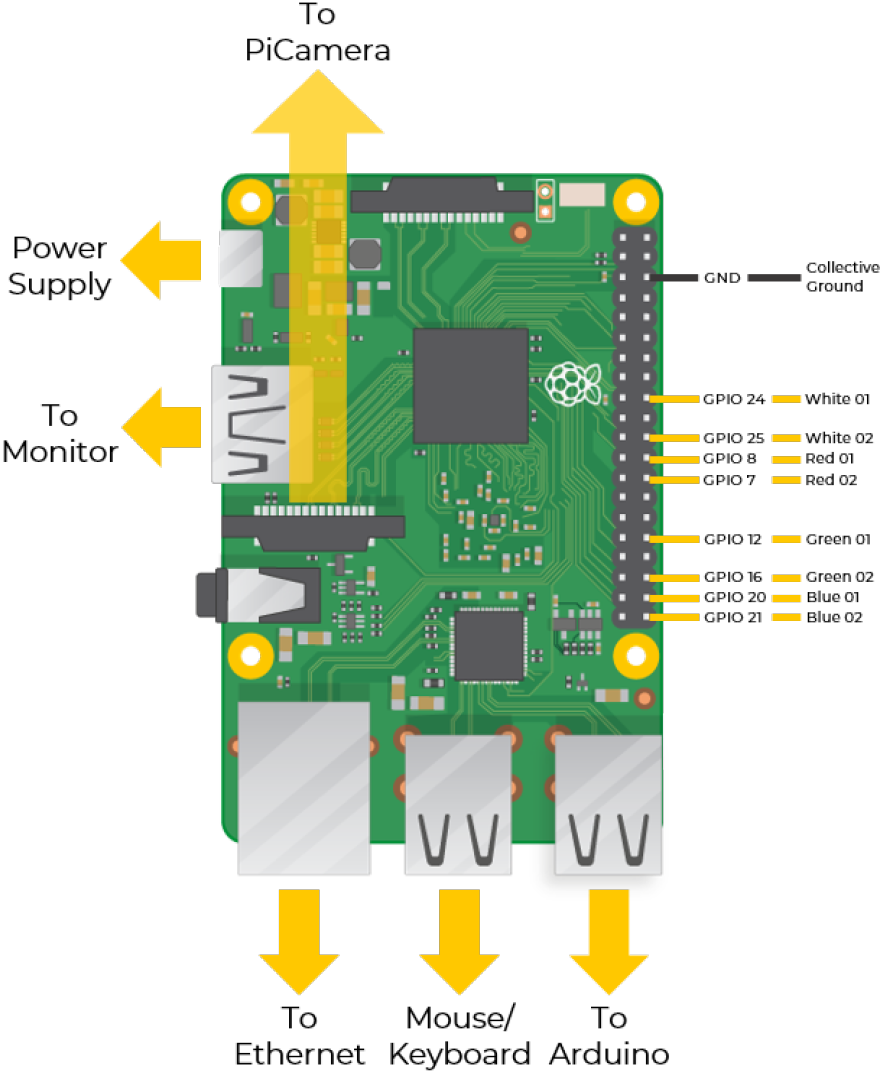
Layout schematic for Raspberry Pi. **Yellow arrows** indicate the location of cables to be attached to the Pi: power supply, monitor, ethernet cable, mouse/keyboard, and Arduino/CNC device. **Yellow lines** indicate path of wiring to from GPIO pinouts (named) to LEDs (named). A single collective ground wire (**black line**) is used to ground all LED wires simultaneously. Raspberry Pi image from (17).

## Incubator Incorporation

1. Detach any cables connecting the *Incubot* to the Pi3B+, AUR3, or power supply. Leave the connections in place on the *Incubot* for ease of re-connecting.
2. Ensure a grating is present on the bottom rung of your incubator and confirm this is securely in place and level.
3. Remove enough gratings above this to facilitate the maximum height of the *Incubot* (Z-axis lead screw).
4. Spray down the disconnected *Incubot* with 70% Ethanol and use an ethanol-soaked wipe to scrub the build as much as possible.
5. Apply WD40 or alternative lubricant to all bearings, moving parts, and parts potentially prone to corrosion to prevent damage from the humidity within the incubator.
6. Re-spray the *Incubot*, and place quickly onto the bottom grating of the incubator. Ensure the *Incubot* is placed with the Y axis motor towards the hinge of the Incubator, to allow for the full range of motion in X, Y, and Z axes.
7. Ensure all wires and cables connected to the *Incubot* are led safely out of the incubator towards the opening side of the incubator. Orientate the X, Y and Z axes so that the Pi Camera is at the furthest possible distance from the opening of the incubator
8. Leaving a little bit of slack in the cables, affix them to the external incubator wall using tape, making sure to spread the cables so that there are no major gaps in the incubator seals that could allow for gas leakage while the doors are closed.
9. Re-connect the cables to their appropriate positions in the Pi3B+ and the AUR3.
10. Return the X, Y, and Z axes to their “resting position”. For our lab, we choose to bring the Y axis as far away from the opening as possible, and the X axis as close to the opening as possible. The Z axis will be left at a height suitable for future imaging, however the level of the Z axis is not important at this point in the construction.

## Software Installation and Setup

1. Install Raspbian onto Raspberry Pi (via Noobs is recommended)
2. Install dependencies (using Python 3.7.3):

a. Kivy
b. NumPy
c. PIL
3. Install Additional Software

a. Arduino IDE
b. GRBL (install directly onto the Arduino, this will allow for cartesian control of the XYZ axes)
c. VNC (or alternative remote screen system compatible with Raspbian)
4. Download all necessary files from the OSF repository onto the Desktop

## Operation Instructions

### Initial Operation

Once the hardware has been established, code imported onto the Pi3B+, and the software is installed correctly, the device is ready to be calibrated. The depth of calibration will depend on the intended use of the *Incubot* and can be update at any time. Calibration focusses on the key steps:

1. Validation of Motion
2. Calibration of Plate and Well Location
3. Objective Lens Calibration
4. Imaging Folder Generation

### Validation of Motion Settings

Included in the files for the *Incubot* is a python script named “MotionValidationProtocol.py”. This script commands the *Incubot* to move in relative motions in positive and negative directions in the X and Y axes, saving images after each movement. The goal is to measure the observed motion compared to the motion requested using G-code. Establish a scale graticule (ideally greater than 10 × 10 mm in size, with graticule markings forming a cross, with graticule markings less than 500 *μ* m apart) within the field of view (FOV) using the “CameraPreview.py” script (via Thonny, a python integrated development environment standard on Raspbian OS), exiting out of the script once the scale bar is located centrally within the FOV, and completely in focus. Leave the system for a few minutes to ensure the temperature within the incubator returns to a normal state to prevent effects of thermal drift on focus. Review the code for the “MotionValidationProtocol.py” script, altering the USB settings if your Arduino is connected through a different port to the one specified in the base c ode. Ensure the script is also sending files t o a n a ppropriate f older for image analysis later. You may wish to alter the camera parameters, or the size of the image to be acquired, this is perfectly acceptable and should not impact the validation process. Run the script using Python3 via the terminal to allow for emergency exit (Esc) from the script. Once the script has completed, the preview window will close, and you will be able to access the files. T here s hould b e 400 images saved within this folder, each labelled according to the axis, repeat number, and whether it is the start image or the end image. Transfer all image files from the appropriate folder to a standard computer/laptop with FIJI (18) installed. Open ImageJ and ensure the program is up to date, updating the program if necessary. Open the script “PairwiseStitchingMotionValidation.ijm” within ImageJ. Alter the “folder” parameter to reflect the folder you have placed your images into. The ImageJ script will sequentially stitch images within this folder, two at a time, recording the pixel translation along with other details of the stitching process. If a results box is currently open, ensure you clear this, then run the stitching script and leave to complete. Once the script has completed, you will be left with a results box with 1600 lines of information. Copy this entire set of information into the first column of the sheet “Raw Data Input” of the template excel file “Motion Validation Template.xlsx”. Columns to the right of this column will then extract the X and Y pixel translations of the stitching process from the text. The first 5 columns after the raw data (B-F) will contain information relating to movement of the X and Y axes in the positive direction. The next 5 columns (G-K) will contain information relating to movement of the X and Y axes in the negative direction. Copy the data in column “F” and paste (Paste>Values) into the cell A2 of sheet “Positive Movement Filtering”. Highlight column A within this sheet and apply a filter (Home>Editing>Sort & Filter>Filter). Press the drop-down triangle and scroll down to the bottom. Uncheck the “Blank” values to hide empty columns. Copy this column and paste (Paste>Values) into the cell A2 of sheet “Positive Movement Collation”. Calculate the number of μm per pixel using an image from your *Incubot* setup of a scale graticule and use the line function to measure the number of pixels in a known distance. Calculate the number of pixels per μm. Input this scale factor into the cell C1 of sheet “Positive Movement Collation” default filled with “PAST CONVERSION FACTOR HERE”. Also input this scale factor into cell C1 of sheet “Negative Movement Collation” for later use. Column B of “Positive Movement Collation” will now fill with the translation distance in *μ* m, converting your original pixel values using your scale factor. The tables to the right will also fill with the appropriate data reflecting the division of the data into the requested motion intervals in each axis. If you have elected to change the requested motion values or alter the number of repeats within the “MotionValidationProtocol.py” script the table will be ineffective and you may have to separate the data manually or adapt the template to suit your requirements. Repeat this process for sheet 4 “Negative Movement Filtering” and sheet “Negative Movement Collation” using the negative movement data from column K of sheet “Raw Data Input”. If the calculated movement is close enough to your requested values for your work, then no further alterations are necessary. If the values in a specific range are insufficient for your studies: 1) check the hardware for appropriate tension across the X and Y axes, confirm alignment of the rails, and confirm no major structural issues are occurring. 2) Re-run the “MotionValidationProtocol.py” script and repeat data analysis steps. If the problem persists, determine the error occurring in each axis, and alter the parameters within GRBL via Arduino IDE to reflect the appropriate number of steps to achieve the movement you are expecting. To do this open the Arduino IDE and open a serial connection to the monitor. Type “$$” and press “enter” to bring up the current settings. Alter your specific setting by typing “$”, the number of the setting, followed by an “=” sign, and the new setting value. Repeat the “MotionValidationProtocol.py” script and confirm your changes were effective. Please note that the “MotionValidationProtocol.py” script performs positive movement and negative movement in sequence, and thus any issues with backlash will be more apparent in the lower range of motion tested. Therefore, it is more reliable to trust the higher end of tested motion to prevent overcompensation and thus over-movement at higher requested distances.

### Calibration of Plate and Well Location

The depth of this calibration will depend on the range of plate-types planned on being used within your work. It is important that validation of motion has been performed extensively and that movement is deemed reliable before moving on to this step. This process will get you to identify key well location landmarks (Fig 10.) to ensure imaging sweeps are accurate to your plate-holding location specific for your system. Establish the Arduino IDE and open a serial monitor connection with the *Incubot*. Place a plate of the type most frequently used (I’d recommend starting with 6-well to familiarise yourself with this process), ensuring the plate is orientated with well A1 closest to the *X* = 0 and *Y* = 0 home position of the *Incubot* optics. Ensure the hardware is in its *X* = 0 and *Y* = 0 positions and that the Z axis is in roughly the correct position. Send the G-Code “G92X0Y0Z0” through the serial connection to establish this position as the home coordinates. Move the X axis and Y axis to roughly appropriate positions using “G90” to ensure absolute movement. Run the script “CameraPreview.py” via Thonny and wait for camera establishment. Plate landmarks referred to from herein are as shown in figure 10. Adjust X and Y coordinate positions until point A1S is visualised. Record this coordinate location. Increase the X axis value until the analogous part of well A2 is visualised (A2S) in the same relative position within the FOV. Record these coordinates. The difference between these X values is your Δ X variable. Confirm this value by moving to the predicted location for A3S based on your Δ X. Return to the initial position for well A1S. Adjust the Y axis coordinate until position A1I is visible. The difference between the Y coordinate for A1S and A1I is your well diameter, it should match with the manufacturer’s specifications for your plate. Now adjust the Y axis coordinates until B1S is visible within the FOV. Record these coordinates. The difference between the Y coordinates of A1S and B1S is your Δ Y value. Record this value. If using a 12-well or higher plate type, confirm this by incrementing the Y axis with the Δ Y value and observe if the analogous portion of well C1 is visible in the same relative position within the FOV. Adjust the X and Y coordinates until you have identified the position A1L. Use the X coordinate of this point, and the Y coordinate of A1S as your starting point for well A1. Calculate this relative position for each of the wells on your plate using the values obtained for Δ X and Δ Y. Depending on the plate-type you are calibrating, select the appropriate function from lines 876-1003 (“IncubotGUI.py”). Adjust the arrays “xCoordinates” and “yCoordinates” to reflect these calculated coordinates. The arrays are read in the order A1, A2, A3… B1, B2, B3… C1… Copy these arrays into the xOrigCoordinates and yOrigCoordinates variables. Once performed, provided the plate location is stable between experiments, this calibration will only need to be performed once for each plate type. If at any point the plate-holder design is rearranged, or altered in any way, this calibration step will need to be re-performed.

**Figure 10.**
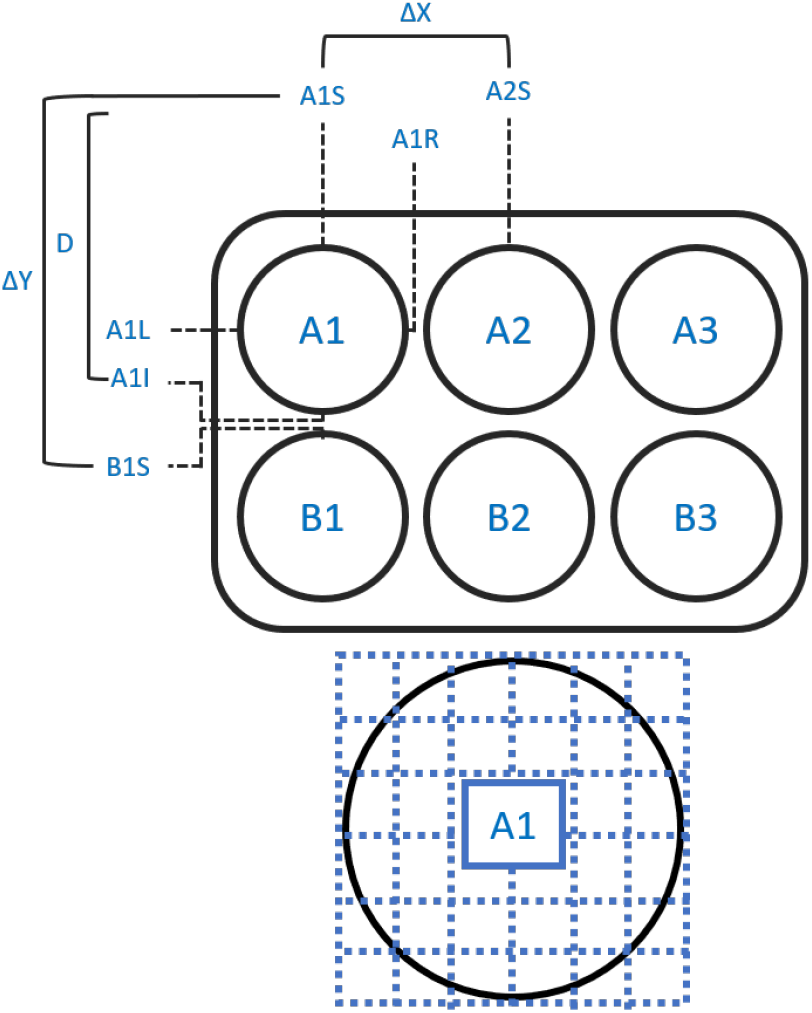
Schematic Representation of a 6-Well Plate with Landmarks for Well Location Annotated. The positional points for well A1 are defined a s the top-most (A1S) the bottom-most (A1I), the left-most (A1L) and the right-most (A1R) points on the well. The difference between A1S and A1I will give the diameter of the well (D), as will the difference between A1L and A1R which should be the same. The difference between the same landmark on adjacent wells will give the distance in X (A1L-A2L or A1R-A2R) between wells (Δ X), and the distance in Y (A1S-B1S or A1I-B1I) between wells (Δ Y). Full available imaging area is shown to the right, with segments for later user selection highlighted.

### Objective Calibration

If using a different objective lens to that described in this build, you will need to adjust the objective lens parameters within the “IncubotGUI.py” script (lines 682-708). Simply attach the objective lens to be used (infinity conjugate is necessary to keep the light path short enough for use, however a modified longer lens tube would facilitate use of a finite conjugate objective lens), and open the script titled “CameraPreview.py”. Adjust the camera parameters to capture and display a preview of the appropriate dimensions for normal *Incubot* functioning, save the script, and run the script using Python3 via the terminal. Establish focus on a graticule with known dimensions and exit out of the script. Un-comment lines 45-47 (“CameraPreview.py”) to enable image acquisition, input desired image save location (line 45) and image name (line 46), save the script, and re-run from the terminal. Transfer the resultant image to a desktop computer and use determine the scale factor (pixels per μm) for the objective lens using FIJI. Use this scale factor to determine the size of the FOV in mm (both X and Y). Multiply these numbers by a factor of 0.8 to give the X and Y increments when using this objective to ensure enough image overlap for image stitching during image processing and analysis. Update the “IncubotGUI.py” script at line 682-708 within the magnification setting functions with the modified values for xMod and yMod, depending on the magnification you are calibrating for. If using a difference objective lens magnification, it would be worth updating the label for one of the provided magnification buttons, along with updating the function to reflect your different objective lens specifications.

## Image Folder Generation

Additional parameters at the start of the “IncubotGUI.py” script specify folders of importance for image saving. It is crucial that you determine and create an appropriate imaging folder for centralised image saving, whether that is on the raspberry pi itself or on an external storage device. Create a folder in your desired location and paste the folder location into the variable “folder” (line 105, “IncubotGUI.py”). If you possess a subscription to a large-volume Dropbox, I would recommend following the instructions within (19) and establish the settings as desired. Rename the Dropbox Upload script “dbupload.py” and uncomment the import bpupload (“IncubotGUI.py”, line 22), and ensure it is in the same directory as “IncubotGUI.py”. Please note that the *Incubot* will upload between imaging sweeps and will wait for upload to complete prior to beginning the next sweep, potentially impacting time between sweeps. A very good internet connection is recommended if Dropbox upload is selected.

### Routine Operation (Fig 11, 12)

An annotated version of the *Incubot* GUI is provided in figure 11. For a quick-start GUI guide, please review figure 12.

**Figure 11.**
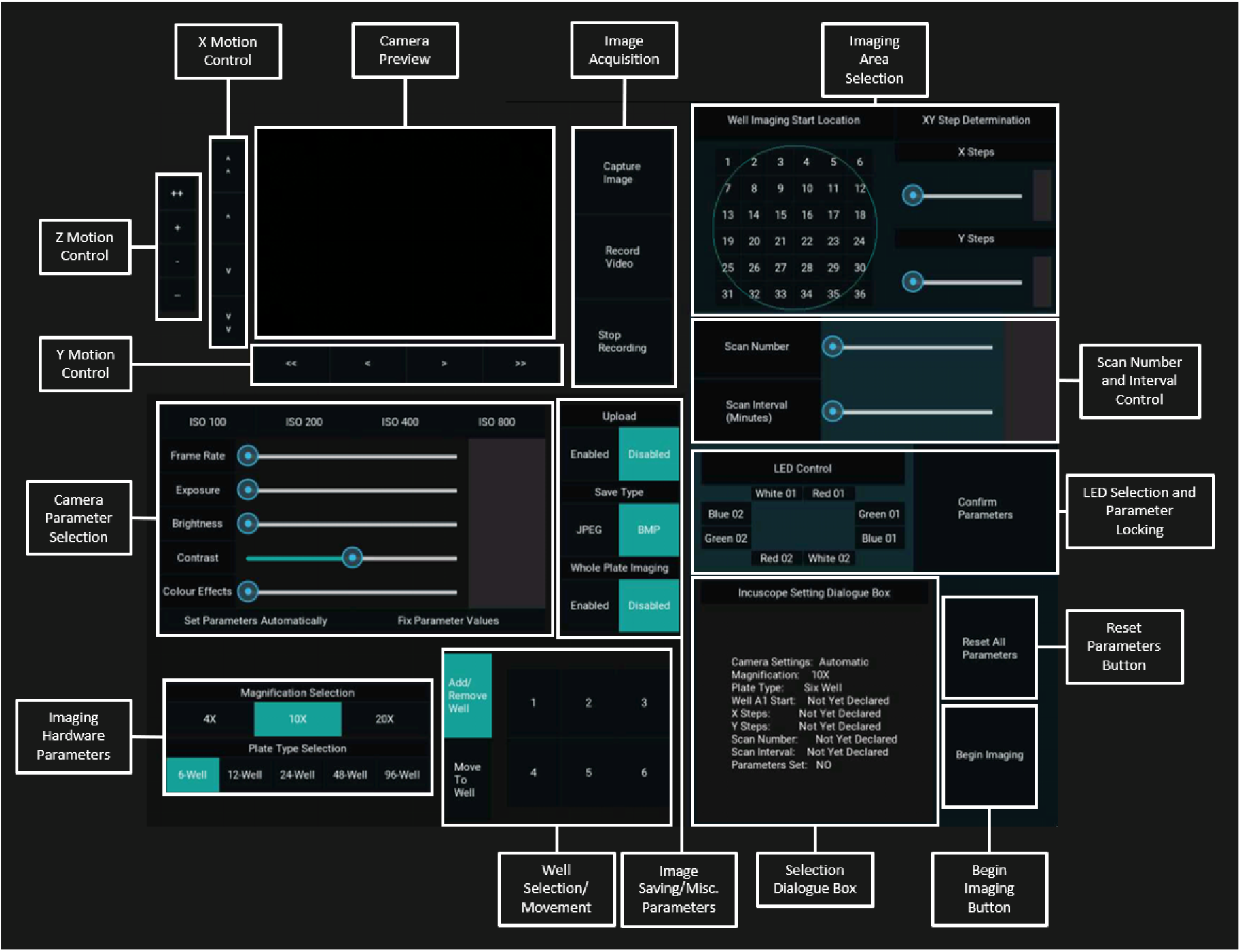
*Incubot* GUI with Key Areas for User Selection Annotated

**Figure 12.**
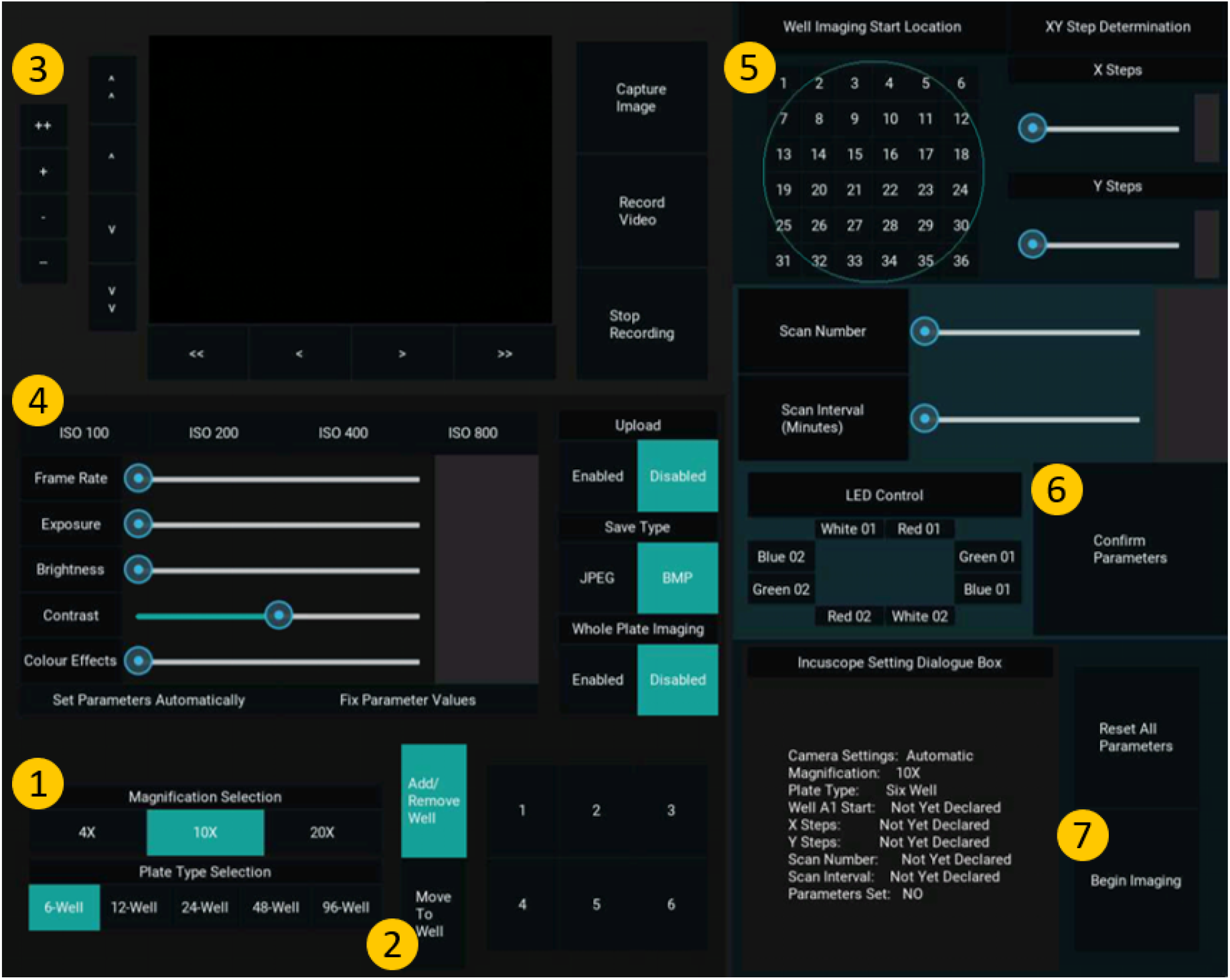
Quick-Start Operational Guide to Routine Use of the *Incubot* GUI. 1) Select your hardware settings (magnification and plate type). 2) Select the “Move to Well” button and select a well of your choice from the grid layout to the right of it. 3) Adjust the X and Y positions until cells are in view and adjust the Z position until cells are in focus. 4) Adjust the image acquisition parameters if needed and alter any image acquisition parameters you wish to. 5) Select an appropriate start location for your experiment and select the number of images in the X and Y axes you wish to collect. Alter the scan number and scan interval parameters to suit your experimental needs. 6) Once all settings are to your liking, confirm the parameters by pressing the “Confirm Parameters” button. 7) Once sure that imaging is established to your liking, press the “Begin Imaging” button to begin the process of imaging. The *Incubot* will be unresponsive during the imaging run, so if emergency stopping is required, the Pi must be physically switched off and back on again. Note that this will result in the loss of positional XYZ information and the *Incubot* will assume the position it is left in is X0Y0Z0. You may need to manually move the *Incubot* to its X0Y0 manually, or via the Arduino IDE once rebooted using negative X and Y coordinates

You may manually determine the appropriate Z level for general imaging by using the “CameraPreview.py” script with a small image preview in conjunction to sending G-code directly using the Arduino IDE program. To do this, establish an image preview and send G-code to move the X and Y axes to an appropriate coordinate for sample visualization. Adjust the Z axis until focus is achieved. Command this location as the new Z0 point using the G-code command “G92Z0”. Alternatively, this can be performed within the *Incubot* GUI. Select your objective lens magnification and plate-type (bottom left corner). The box to the right of this section will update to reflect the plate type you’ve selected. Press the “Move to Well” button to use the simulated plate to move to the default XY coordinate specified for the selected well, which will not have the well base visible. Use the X and Y motion control sections to move the FOV to a region within the well where cells should be visible. Use the Z motion control section to adjust Z axis until focus is achieved. It may be necessary to adjust your Camera Parameter Selection options to optimise image preview/acquisition parameters. The LED “White 02” will be active by default, so you may also need to select alternative LED(s) using the LED Control Box (middle right). Select/deselect options relating to Dropbox uploading, image file save type, and whole-plate imaging using the buttons within the Image Saving/Misc. Parameters section. If you would like to acquire an image or recording now for your records, you can do so using the Image Acquisition section to the right of the Camera Preview. If you wish to exclude certain wells from imaging, press the Add/Remove Well button and then select the wells you wish to exclude (excluded wells will be highlighted in dark blue).

To alter parameters relating to your imaging experiment needs, use the right side of the GUI. The imaging area by default will start outside of each well, at the minimum X and Y coordinate location as defined b y t he u ser d uring plate and well calibration. To alter this start location, select an appropriate button within the Well Imaging Start Location layout. A representative well is shown using a blue circle to allow for visualization of your rough start location. Select the area you would like to image using the XY Step Determination section to select your X and Y imaging dimensions in terms of images along the X axis and images along the Y axis. Below this, select the number of times you would like to scan the plate, along with your desired interval between scans (start time to start time). It is possible to set impossible requirements here by requesting an interval that is shorter than the actual time required for a single scan. Under these conditions, the *Incubot* will begin scans immediately after the previous one, however it will not inform you if this is expected to be the case. You can reduce the time required for a complete scan by reducing the area of imaging, or by saving in JPEG format as opposed to BMP. Ensure the LED(s) you would like to use during imaging are selected in the LED Control section (selected LEDs will be highlighted in blue). Once you are happy with your parameter selection, set the parameters using the Confirm Parameters button.

At this point you may wish to review your settings selection by observing the *Incubot* Setting Dialogue Box. If you wish to change parameters you may either alter the sliders/button individually or press the Reset All Parameters button to begin the process from scratch (this will only affect parameters, and not the current *Incubot* coordinates). Once all parameters are to your liking, you are ready to begin imaging by pressing the Begin Imaging button. This will trigger the start of the scanning process. The *Incubot* will begin imaging at the first non-excluded well in the order of A1, A2, A3… B1, B2… etc. Once imaging has concluded, provided you have opted not to automatically upload your files to Dropbox, exit out of the GUI by pressing the Esc button on your keyboard. The imaging folder will be within your selected directory ready for transfer to a desktop computer for image processing, stitching, and analysis.

### Routine Maintenance

Due to the use of the *Incubot* within a tissue culture incubator, regular coating of parts with WD-40 (or alternative lubricant) is required to prevent corrosion of parts. I recommend performing this every 2 months, but more frequently is better.

Cleaning should be carried out when needed by removing the *Incubot* from the incubator and scrubbing down the build using tissue soaked in ethanol. WD-40 should be reapplied following each cleaning. Provided there are no tissue culture infections or visible build-up of matter, cleaning of the build can be kept to roughly every 6 months.

## Validation and Characterisation

### Hardware Validation

#### Plate Stability

Plate stability was assessed by imaging a single location of a USAF 1951 resolution testing slide (21) every minute for 60 minutes (“StationaryStabilityTesting.py”). The coordinate location of a defined high-contrast line corner within the image was manually determined, and the displacement over time was calculated in both the X and Y axes. The X axis (Fig 13A.) showed a higher level of stability than the Y axis (Fig 13B.), and only moved a on average ± 1.3 *μm* over the time span. The Y axis showed greater displacement (±5.8 *μm*), indicating a minor degree of motion between images, however as the variations showed no trend over time (linear regression, P=0.1998), there is no evidence of persistent drift.

**Figure 13.**
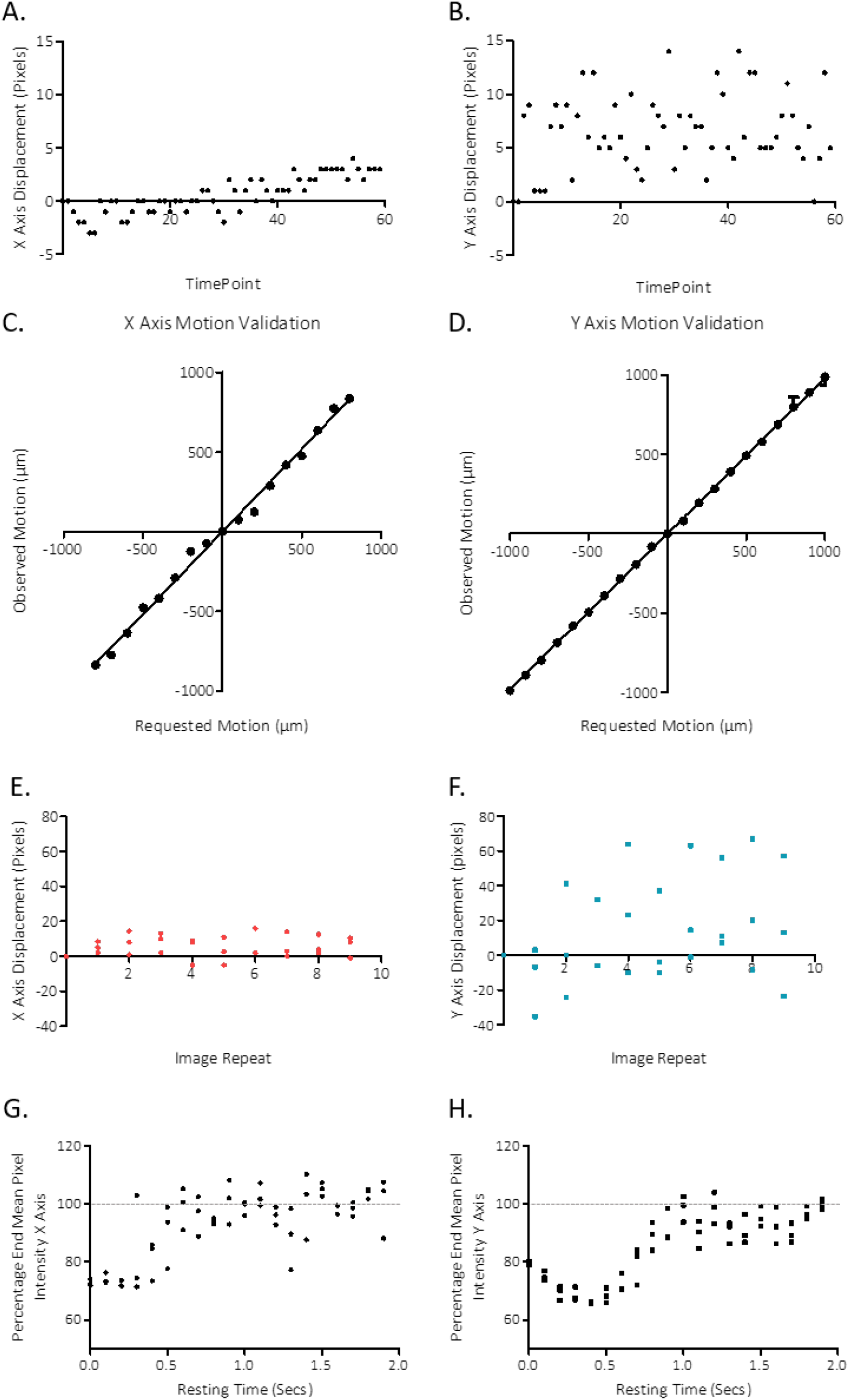
Stability and Motion Validation of the *Incubot*. A) Deviation of an X coordinate of a landmark within an image over 60 minutes of imaging. B) Deviation of the Y coordinate of a landmark within an image sequence over 60 minutes of imaging. C) Distance moved by the *Incubot* in the X axis compared to requested distance via G-Code. D) Distance moved by the *Incubot* in the Y axis compared to requested distance via G-Code. E) Deviation in X coordinate location of a landmark when the *Incubot* is commanded to move from a set location from a different location. F) Deviation in Y coordinate location of a landmark when the *Incubot* is commanded to move from a set location from a different location. G) Sharpness (20) of images taken by the *Incubot* following movement in the X axis with variable rest time delay durations. H) Sharpness (20) of images taken by the *Incubot* following movement in the Y axis with variable rest time delay durations.

#### Movement Fidelity

Movement fidelity was calculated according to the protocol established earlier in this manuscript (Operation Instructions > Validation of Motion Settings). Motion was requested in the range of 0.1-1.0 mm (Y axis) or 0.1-0.8 mm (X axis) in mm increments. Following optimisation of the GRBL parameters controlling steps per mm, motion was highly reliable for a given requested motion, showing very little variability. Observed motion was sufficiently comparable to requested motion in both X and Y axes for our purposes (Fig 13C,D.).

The reliability of coordinate motion was also calculated by repeated imaging of a fixed tissue culture plate. The *Incubot* was commanded to scan single points within wells 10 times, moving back to the home location between each imaging sweep. Location of a structurally defined point (cell process termination) was manually determined and the displacement was compared between imaging sweeps for 3 wells in both X and Y axes (Fig 13 E, F.). Y axis repeatability showed more variation than the X axis, however the displacement is not significant enough to disrupt normal image processing/stitching procedures.

#### Minimum Time Between Movement and Imaging

As the *Incubot* hardware rests within a tissue culture incubator, it is impossible to completely protect against vibration and its effect on image quality. The major source of vibration within the incubator comes from the *Incubot* itself during periods of X or Y axis movement, necessitating a rest period between movement and image acquisition. In order to calculate the minimum delay required between movement and imaging, testing was performed to identify the shortest necessary period between movement and imaging. The *Incubot* was set up focused on human dermal fibroblasts (HDF) fixed with 4% paraformaldehyde. We used a script (“MotionRestTesting.py”) to trigger movement of the optics 1 mm in either the X or Y axis and collected images at 0.1 second intervals over a period of 2 seconds. Stability was measured by the sharpness of the image, calculated using the ImageJ plugin “Microscope Focus Plugin” (20).Stability was achieved 0.6 seconds after movement command in the X axis, and 1.0 seconds after movement command in the Y axis (Fig 13G,H.).

### Optical Validation

#### Resolution Testing

Maximum image resolution was determined by imaging a USAF 1951 resolving power grid (Fig 14A.) (21) and also calculating the slant edge modulation transfer function (MTF) in FIJI (22). The FOV was split into 20 subregions in a 4×5 grid, and the test pattern was imaged (using transmitted light illumination) with the smallest resolvable elements moved moved between each of the subregions. The smallest lines clearly visible (*can you also give this as group and element?*)in each quadrant was recorded and converted into its equivalent *lines · mm*^−1^. The majority (16/20) of the quadrants achieved an observed resolving power of 228.1 *lines · mm*^−1^, while 4/20 of the quadrants achieved the slightly lower resolving power of 203.2 *lines · mm*^−1^ (Fig 14B.). Slant edge modulation transfer function (MTF) was also assessed on an image of the USAF 1951 slide taken by the Incubot using transmission illumination at three different pixel resolutions: maximum PiCamera V2 resolution (3280×2464), the default Incubot resolution (1680×1200), and a 4X binned resolution (820×616). The MTF curves are presented in Fig 14,C,D,E. Sensor resolution did not seem to impact the MTF output, and thus the limiting factor for resolution is likely within the optical system.

**Figure 14.**
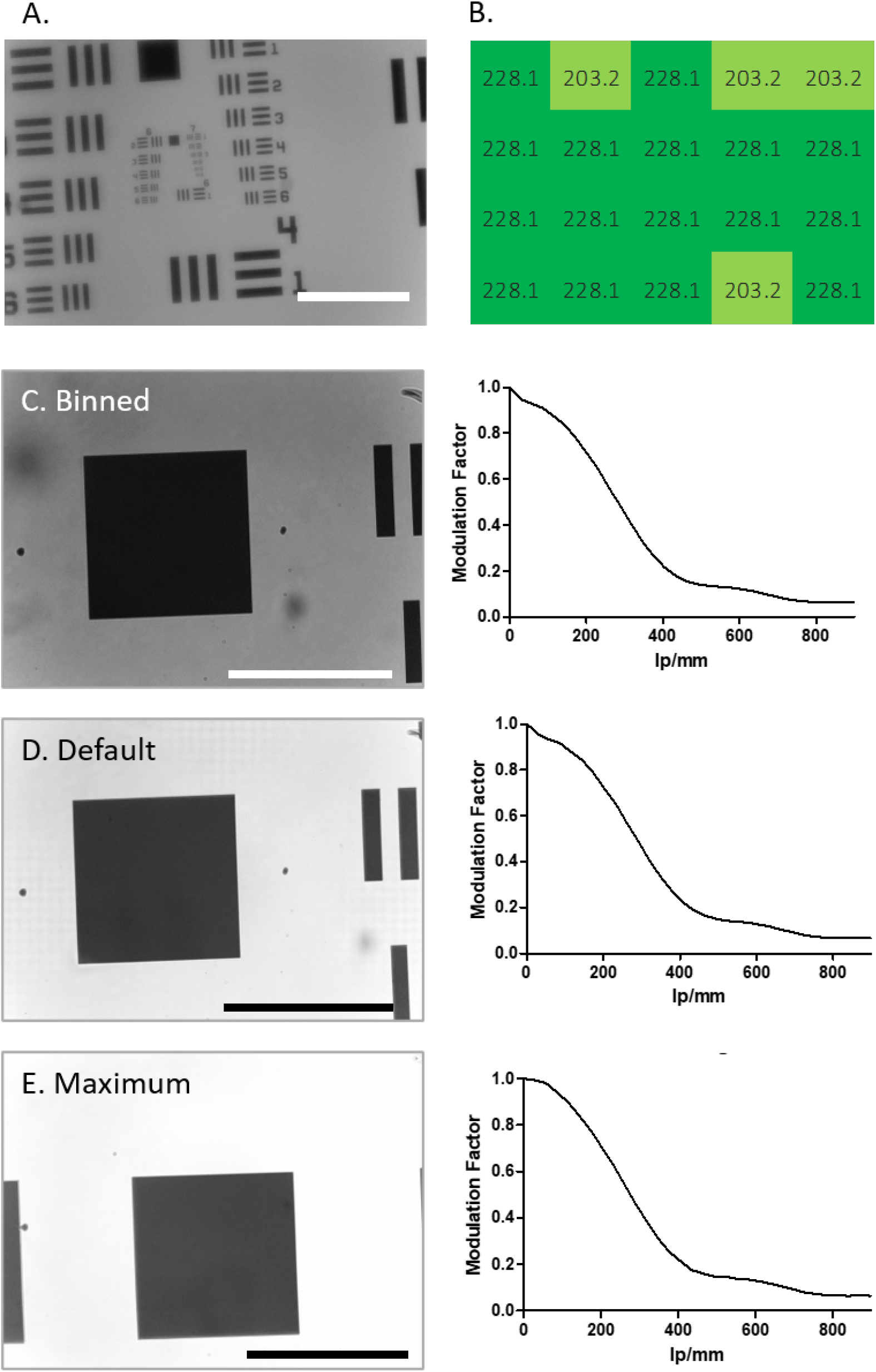
Optical Validation of the *Incubot*. A) USAF 1951 resolution testing grid representative image taken on the *Incubot* at 10X magnification using transmission illumination. B) Maximum resolution (line pairs.mm-1) observed in each quadrant of the *Incubot* field of view (1680×1200 pixels) using USAF 1951 testing grid. Analysis of resolution using slant edge MTF with comparisons between a highly binned image (C), the resolution used for *Incubot* function (D), and the maximum resolution possible on the raspberry picamera optics (E). Slant edge MTF function (22) was applied to the major right-hand slope present in each of these images, with resultant line pairs per mm calculations graphed, and displayed next to each tested setting. Modulation factor is labelled on the y-axes, while spatial resolution measured in line pairs per millimetre (lp/mm) is labelled on the x-axis.

### Validation of Imaging with Multiple Light Sources

#### Reflected and Oblique Illumination

HDF cells were cultured in a 6-well plate and imaged using both white LEDs, then just a single white LED to compare reflected w hite i llumination w ith o blique white illumination. Oblique illumination revealed additional cellular details not visible with dual white LED illumination, highlighting the utility of low-cost single LED illumination at a 45° angle for visualising cellular morphology (Fig 15A.).

**Figure 15.**
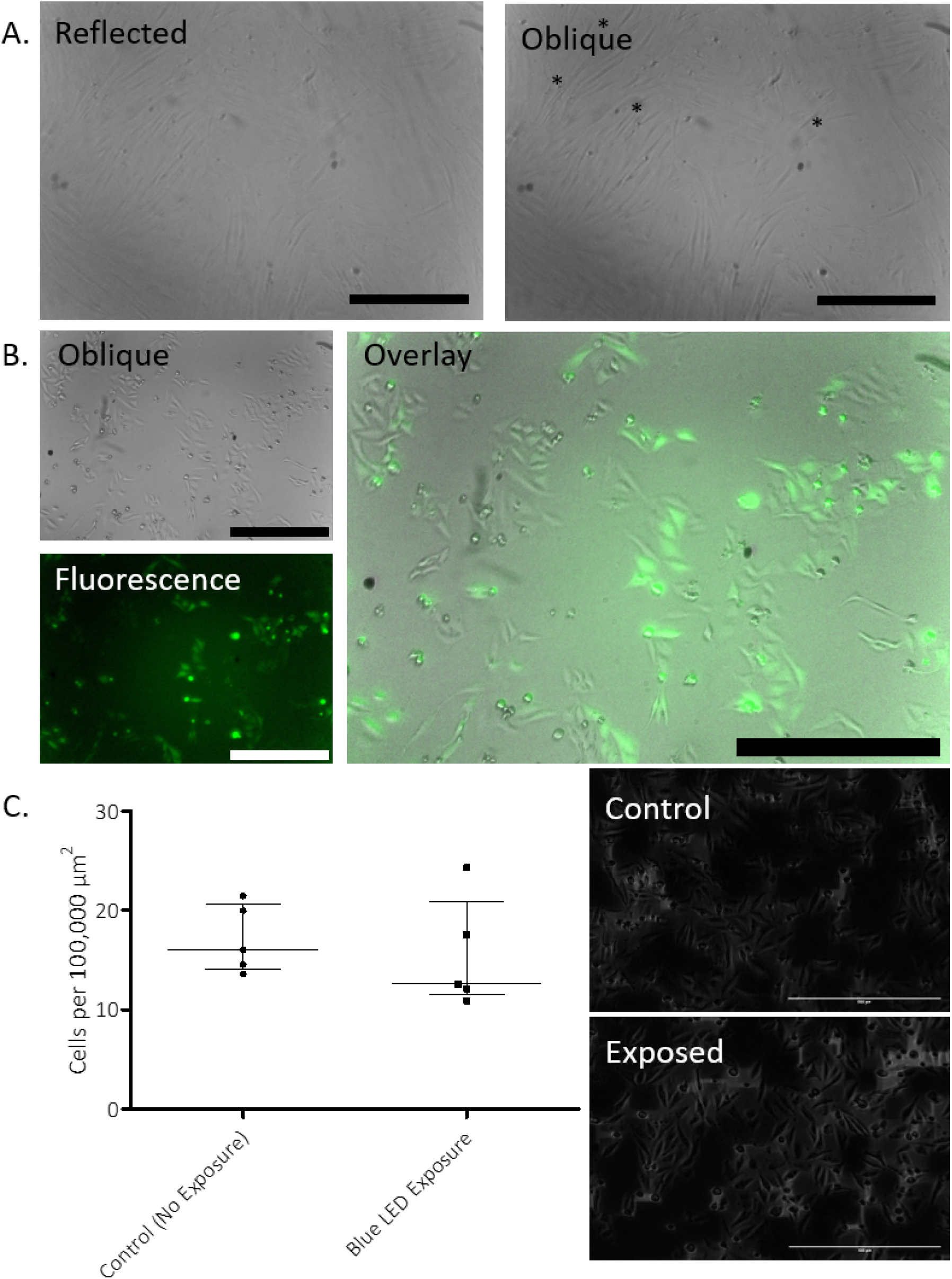
Image Acquisition Validation of the *Incubot*. A) Representative image of live HDF cells using dual white LED illumination (reflected) or single white LED illumination (oblique) with processes uniquely identified using oblique illumination highlighted with (*) (scale bars = 400 μm). B) Live HeLa-GFP visualization using oblique illumination, blue LED fluorescent illumination, and overlay (scale bars = 500 μm). C) Assessment of phototoxicity revealed no significant difference in cell density between LED-exposed and control cell after 24 hours of repeat illumination of live HeLa-GFP cells (scale bars = 500 μm).

#### Fluorescence

For fluorescence imaging the *Incubot* needs to be capable of fluorescence detection, but importantly because t his system needs to be capable of fluorescence imaging repeatedly over a long time frame, it should not induce significant photodamage to cells over the course of this imaging. We assessed this by culturing commercially available HeLa cells stably expressing GFP (AMS Biotechnology). GFP was excited by both blue LEDs and the emission was detected and overlaid well with oblique images taken in the same coordinate location immediately after (Fig 15B.). To assess phototoxicity induced by repeated imaging, two 6-well plates of HeLa cells were seeded at a density of 15,000 cells per well and left for 4 hours to allow for cell attachment. One plate was covered from light within the incubator, while the other was subjected to repeated imaging under blue LED excitation every hour for 24 hours. Following the imaging protocol, the light-protected plate was imaged once in the same relative locations as the imaged plate. Cell density was calculated at this 24-hour point for all wells exposed to the LEDs (fluorescent) and all wells protected from light (negative control) (Fig 15C.). There was no significant difference between the density of cells in wells exposed to the blue LEDs compared to control wells (Mann Whitney, P=0.4206), indicating no significant impact of LED exposure on cell viability.

### Software Validation

#### Raspberry Pi Compatibility

The *Incubot* GUI (“IncubotGUI.py”) was designed to be used on a Raspberry Pi 3 B+ with Python 3.7.3. To date, no alternative equipment has been tested for compatibility with the *Incubot*, however it is theoretically possible to generate a new code for an alternative system.

The initial portion of python code is responsible for importing all the relevant dependencies for running of the full code, this is partitioned into a general section and a Kivy section. Kivy is a GUI designing module for python 3 which allows for generation of “apps” within a python environment (23). Each button within the app is generated within Kivy and bound to a function. The functions vary based on the button, for example pressing the “6-Well” button will allow the raspberry pi to change the variables associated with plate type, such as number of wells, well diameter, and the [X,Y] coordinate locations for the default initial imaging of each well. This button, along with others, also commands Kivy to re-initialise the user interface, allowing for an updated screen view for greater ease of use.

#### Z-Axis Automated Focussing

Z-axis automated focussing was optimised using an image stack taken of low-density HeLa-GFP cells grown in monolayer conditions. A Z-stack with slice intervals of mm was acquired using the *Incubot* (1680×1200, BMP format). Images were assessed for sharpness of edges using a Laplacian filter with, or without, a Gaussian filter or varying radius applied beforehand. The images were manually assessed for the slice with optimal focus, and then the output sharpness measurements were compared to determine which parameters allowed for correct selection of the most in-focus slice. Pure Laplacian filtering, and Laplacian filtering with a Gaussian pre-filtering (radius=1) resulted in the selection of the correct image slice. Increasing radius of the Gaussian pre-filtering step (radius=2, radius=3) resulted in increasingly inaccurate assessment of optimal image slice. Thus, a Laplacian filter was applied to images with a Gaussian pre-filtering step using a radius of 1 pixel.

## Concluding Remarks

The *Incubot* represents an addition to the current open source microscopy options available to research scientists. It has proven ability in white light and fluorescent imaging of live and fixed cell culture monolayer within a tissue culture incubator, maintaining stability and reliability. The low-cost of the build compared to commercial options and the open source nature of the microscope facilitate its accessibility to a wide range of potential users.

## Capabilities

1. Long-term fluorescent or oblique imaging of samples
2. Structurally stable within the incubator
3. Allows for physiological imaging of live cell monolayers
4. User-friendly GUI for easy-to-use microscopy system
5. Individual cells and cell processes visible over time

## Limitations

1. Upload to Dropbox cannot occur simultaneously with imaging
2. Time for imaging may impede on some desired imaging experiments, requiring compromise from the researchers
3. Unable to exit imaging experiment without hard shut-down

## Supporting information

Supplemental Parts List

## Declaration of Interest

The authors declare no competing interests

## Human and Animal Rights

Human and animal rights: none

## Acknowledgements

Thanks must be given to Kevin McCarthy for his help with hardware stability.

## Funding

This work was funded in full by University College Dublin, School of Medicine.

## Author Contributions

GOTM was responsible for designing and testing the *Incubot*,and manuscript preparation. BL and CK were involved in initial design steps of the XYZ translation and the optics systems respectively. ERG provided useful input into the hardware design and stability, along with manuscript preparation. NB was involved in carrying out the resolution testing presented here. MP oversaw all aspects of the project and manuscript preparation.

## Notes

https://osf.io/es3hr/

### Competing Interest Statement

The authors have declared no competing interest.

### Summary of Updates

Figure 5 has been replaced with the correct intended figure.

